# The unrealized potential of herbaria for global change biology

**DOI:** 10.1101/218776

**Authors:** Emily K. Meineke, Charles C. Davis, T. Jonathan Davies

## Abstract

Plant and fungal specimens in herbaria are becoming primary resources for investigating how plant phenology and geographic distributions shift with climate change, greatly expanding inferences across spatial, temporal, and phylogenetic dimensions. However, these specimens contain a wealth of additional data—including nutrients, defensive compounds, herbivore damage, disease lesions, and signatures of physiological processes—that capture ecological and evolutionary responses to the Anthropocene but which are less frequently utilized. Here, we outline the diversity of herbarium data, global change topics to which they have been applied, and new hypotheses they could inform. We find that herbarium data have been used extensively to study impacts of climate change and invasive species, but that such data are less commonly used to address other drivers of biodiversity loss, including habitat conversion, pollution, and overexploitation. In addition, we note that fungal specimens are under-explored relative to vascular plants. To facilitate broader application of plant and fungal specimens in global change research, we outline the limitations of these data and modern sampling and statistical tools that may be applied to surmount challenges they present. Using a case study of insect herbivory, we illustrate how novel herbarium data may be employed to test hypotheses for which few data exist, despite potentially large biases. With the goal of positioning herbaria as hubs for global change research, we suggest future research directions and curation priorities.

## INTRODUCTION

A key challenge for biologists today is to determine how species are responding to the major drivers of global change and biodiversity loss: habitat conversion and degradation, climate change, invasive species, pollution, and overexploitation (Millennium Ecosystem Assessment 2005). Over the past decades, field observations and experiments have informed much of our understanding of biological responses to these major drivers, particularly climate change. However, like all scientific approaches, they have limitations. Experiments are almost always at smaller spatial scales than inferences (e.g., Pelini *et al.*, 2011) and field observations are often restricted to temperate biomes (Wolkovich *et al.*, 2012). Experiments and observations typically only allow researchers to test hypotheses about a single driver of global change, while organisms are generally exposed to many. Perhaps most critically, the majority of experiments and field observations are short-term. Experiments addressing global change are commonly conducted for a few years at most (e.g., Diamond *et al.*, 2012), limited by grant timelines and funding cycles. Field observations are often designed to span latitude and altitude as a proxy for warming, relying on the assumption that patterns across space will represent future patterns across time (e.g., Kozlov *et al.*, 2013). While long-term field observations can span over a century in cases where people across generations collect the same observations, these data are rare and only available for a few phenomena (e.g., Keeling & Whorf 2005; Aono & Kazui 2008). Scientists have increasingly turned to biological collections to expand data across time, space, and taxonomy, thus better matching the scales at which recent global change is occurring (Pyke & Ehrlich 2010). In particular, herbarium specimens, preserved (often pressed) plants and fungi, have been the subject of a new wave of global change research.

The potential of herbarium data for global change biology stems in large part from its temporal extent. Since the 1700s, scientists, including Linnaeus and Darwin, have collected herbarium specimens to describe new species, aid taxonomic classifications, and as part of regional floristic treatments (e.g., Moffett 2014). Although collecting has slowed in recent years in many localities (Meyer *et al*., 2016; e.g., Fig. 1a), scientists and amateurs continue to collect. In many parts of the world, such as the northeastern US, the density of sampling extends far beyond what is available from observations and experiments (Fig. 1b) and encompasses most lineages of vascular plants (Fig. 1c), fungi, diatoms, and groups variously classified as algae. Current estimates indicate that herbaria house over 350,000,000 specimens (Thiers 2016), representing increasingly connected national (e.g., the Consortia of US Herbaria, Chinese Virtual Herbarium [http://www.cvh.ac.cn/news/8]) and international networks of data (Fig. 1d; e.g., Australasian Virtual Herbarium [https://avh.chah.org.au/], iDigBio [https://www.idigbio.org/], and GBIF [https://www.gbif.org/]).

**Fig. 1.**
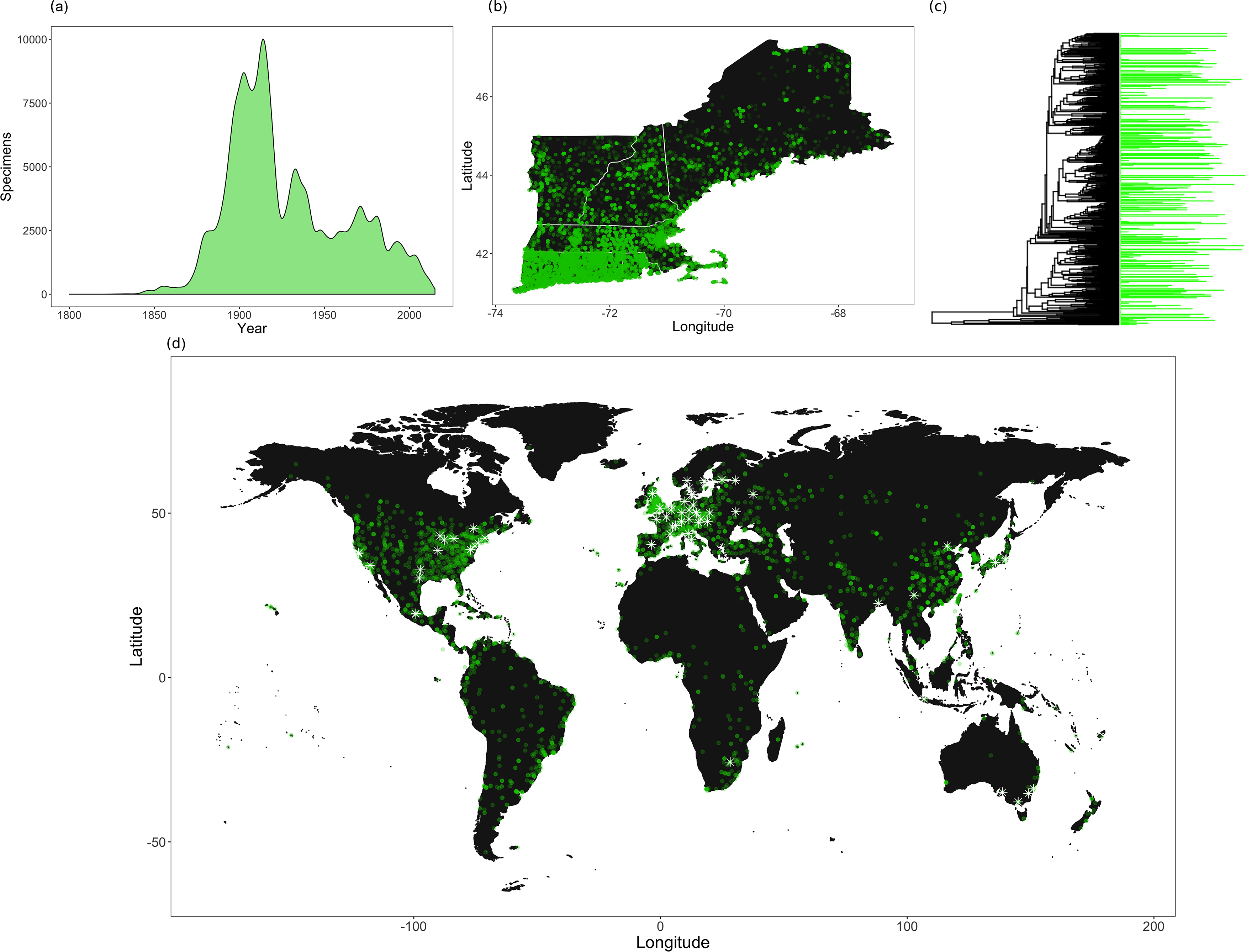
The spatial, temporal, and phylogenetic extents of herbarium specimens. Here we explore the richness of herbarium data, with emphasis on the major herbaria in and area within the northeastern USA referred to as New England (http://neherbaria.org/). (a) The herbaria of New England house more than half a million regional herbarium specimens collected as early as the 1800s, before industrialization and globalization, that can serve as baselines prior to anthropogenic change. The density plot represents the number of collections each year. (b) These specimens have spatial coverage across the region (shown here), and millions more specimens are available from other parts of the world. Though there is clear spatial bias, the spatial coverage exceeds that which is typically available in observations and experiments. (c) These specimens span most branches of the vascular plant tree of life. Here, bars represent log abundance of specimens in families, which are organized by evolutionary history (Harris & Davies 2016). (d) Herbaria are distributed worldwide. Many of the larger herbaria have amassed similar coverage to the New England herbaria across these axes. Here, symbols represent herbaria. Herbaria with more than one million specimens are indicated by white stars.

Scientists today employ these specimens for purposes that could not have been imagined by their collectors. Evolutionary biologists extract and sequence ancient DNA from herbarium specimens to reconstruct phylogenetic relationships or infer population dynamics (Gugerli *et al.*, 2005; Wandeler *et al.*, 2007) and have documented selection on species traits through changes in plant morphology (Kavanagh *et al.*, 2011; Burns *et al.*, 2012). More recently, ecologists routinely use herbarium specimens as occurrence records for determining species distributions (Graham *et al.*, 2004; Newbold 2010) and as records of leafing, flowering, and fruiting phenology (Miller-Rushing et al., 2006; Primack *et al.*, 2004; Bolmgren & Lonnberg 2005; Everill *et al.*, 2014) to understand how these aspects of biology are influenced by rising global temperatures. The value of such data is now well recognized (Lavoie & Lachance 2006; Pyke & Ehrlich 2010; Johnson *et al.*, 2011; Lavoie 2013; Vellend *et al.*, 2013; Willis *et al.*, 2017a). The unrealized potential of herbaria is perhaps most apparent in the opportunities presented by other types of data that are only rarely extracted from specimens but which provide opportunities to assess the effects of global change, including signatures of pollination efficiency, pollution concentrations, physiological characteristics, nutrient concentrations, pathogen loads, morphological and anatomical traits, genotypes, endophytes, and herbivory (Table 1). These could allow researchers to address hypotheses about species and ecosystem responses to global change, from shifts at the species level to changing environmental processes, including nutrient cycles, changes in air quality, and biological control (Table 1).

**Table 1.**
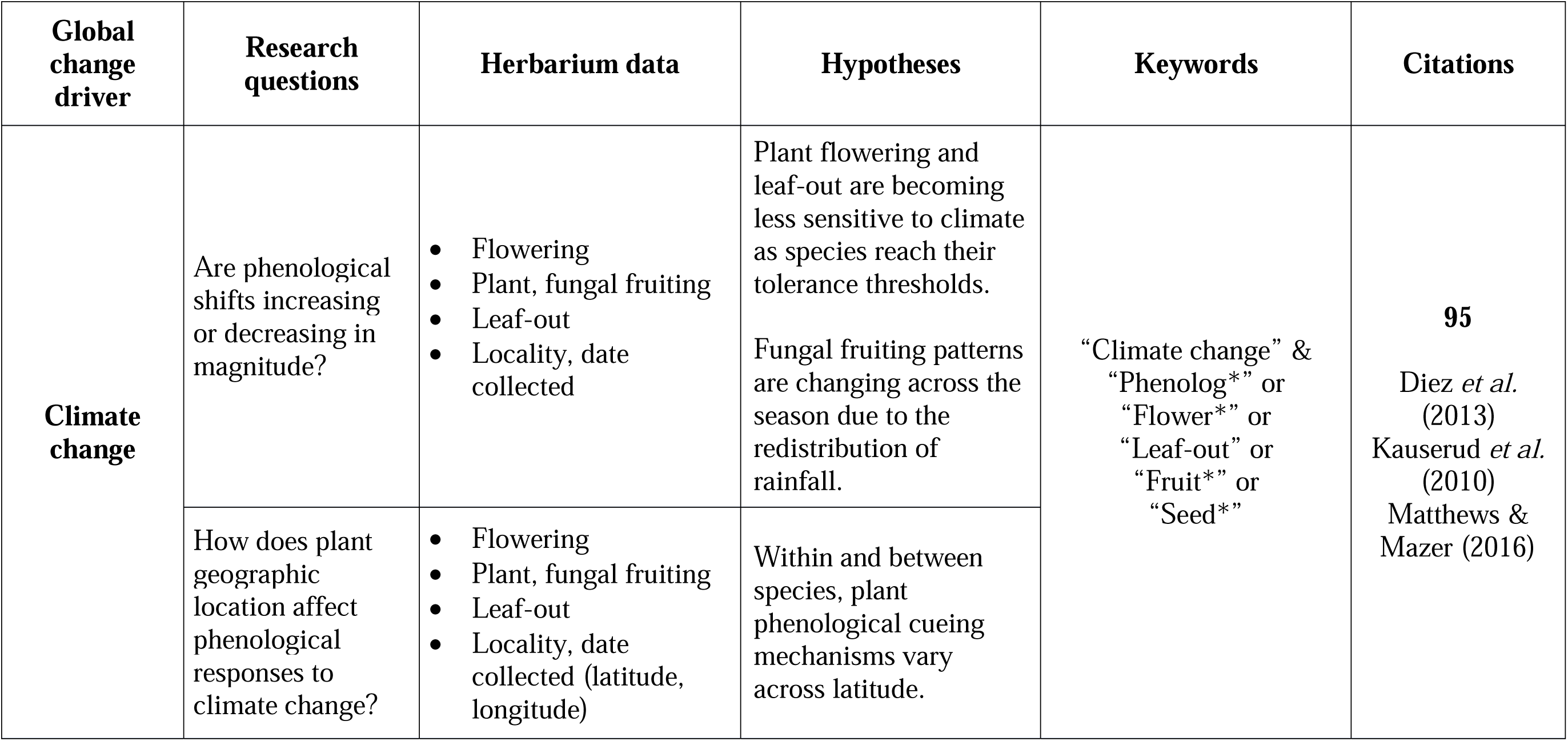
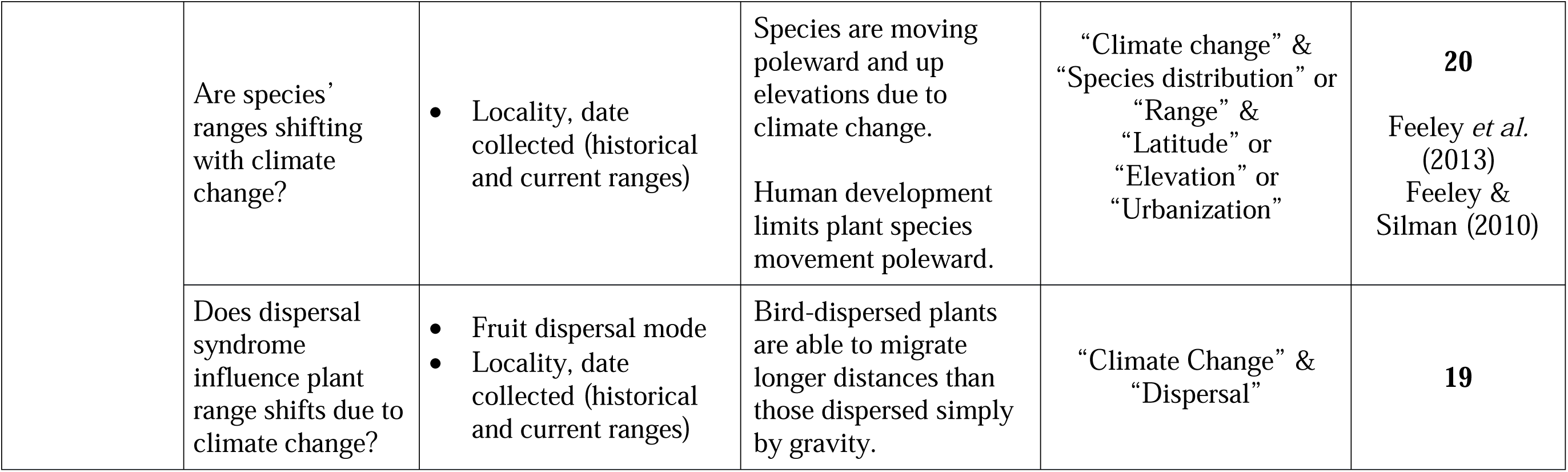
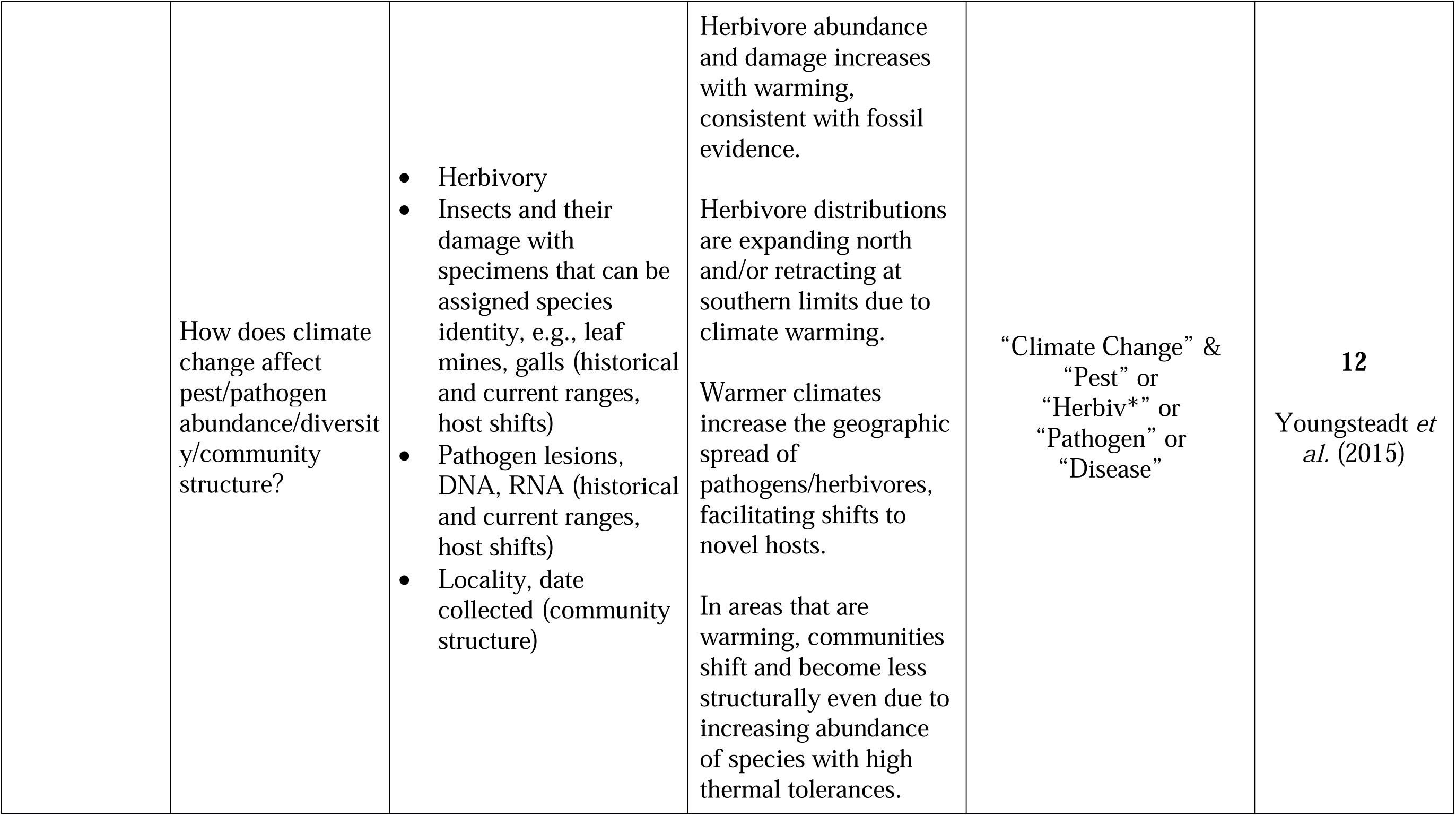
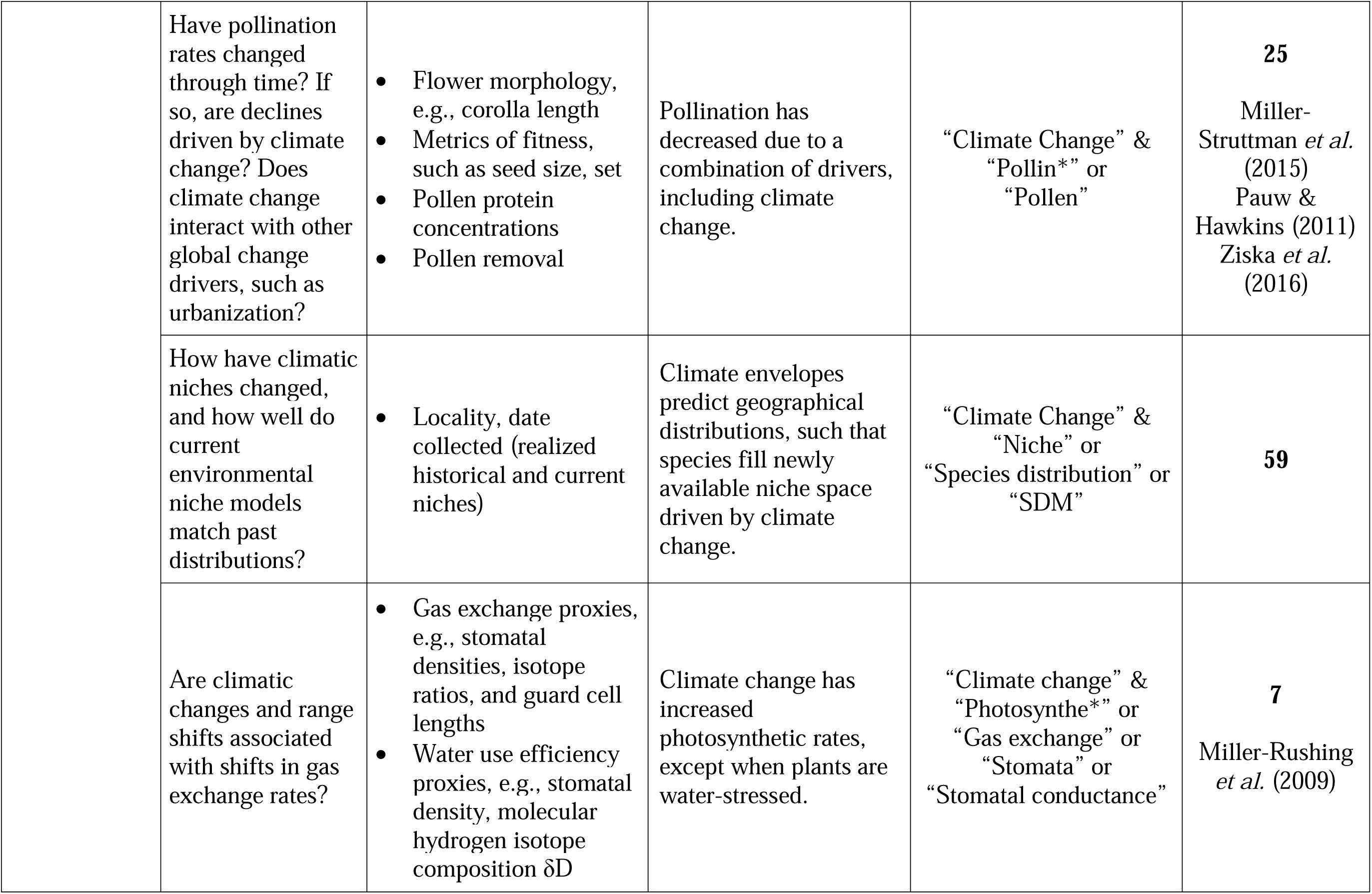
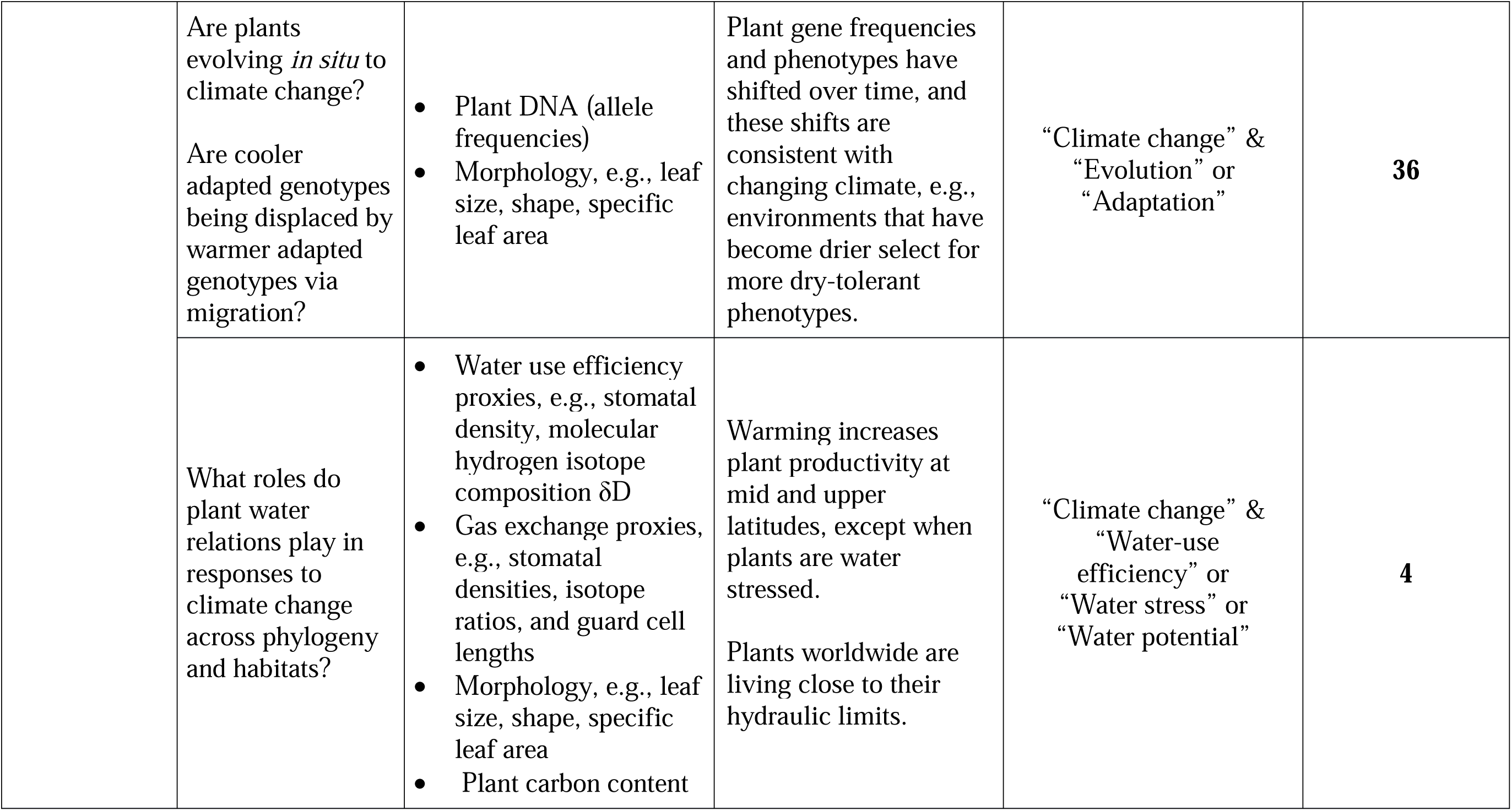
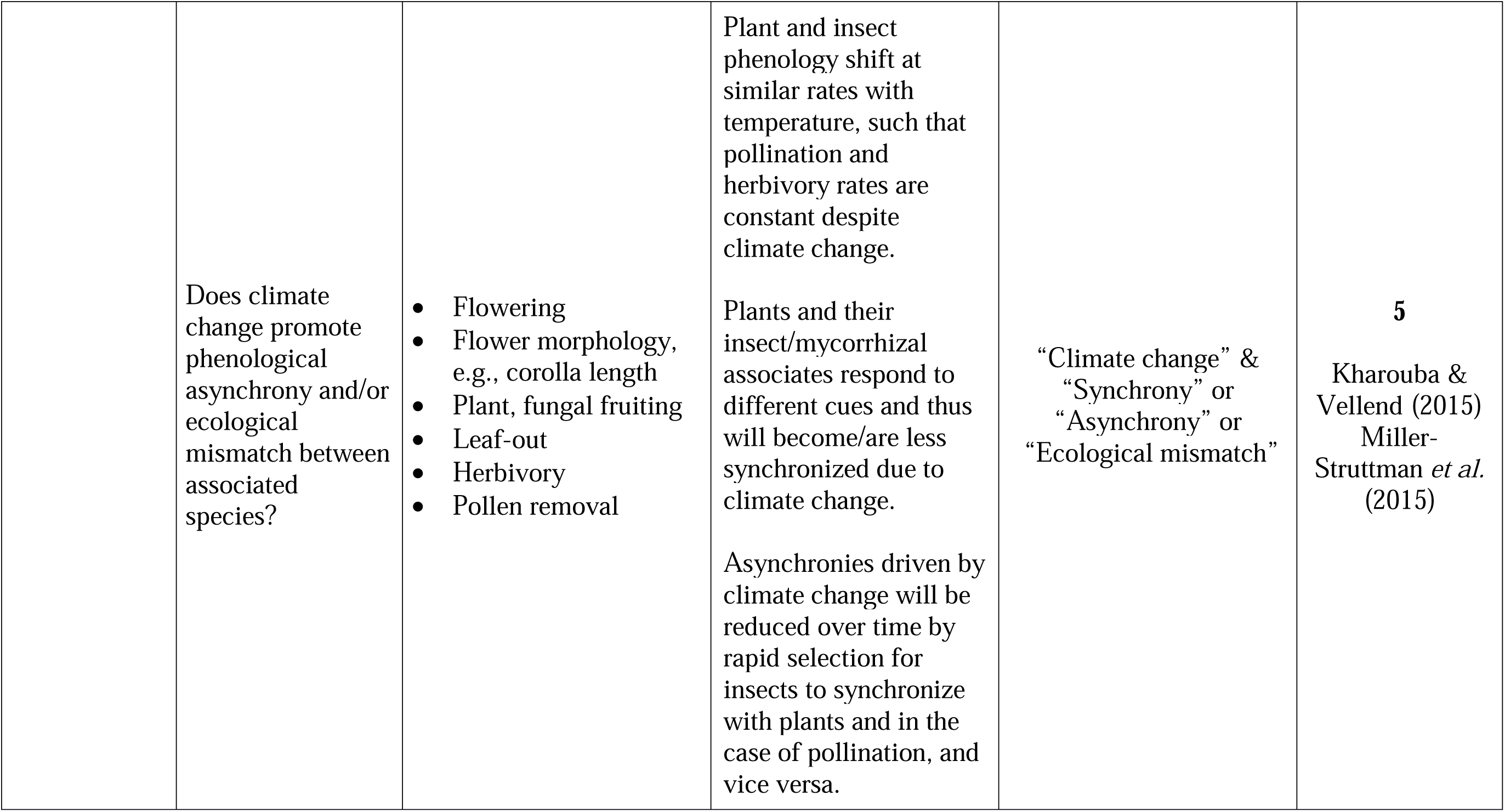
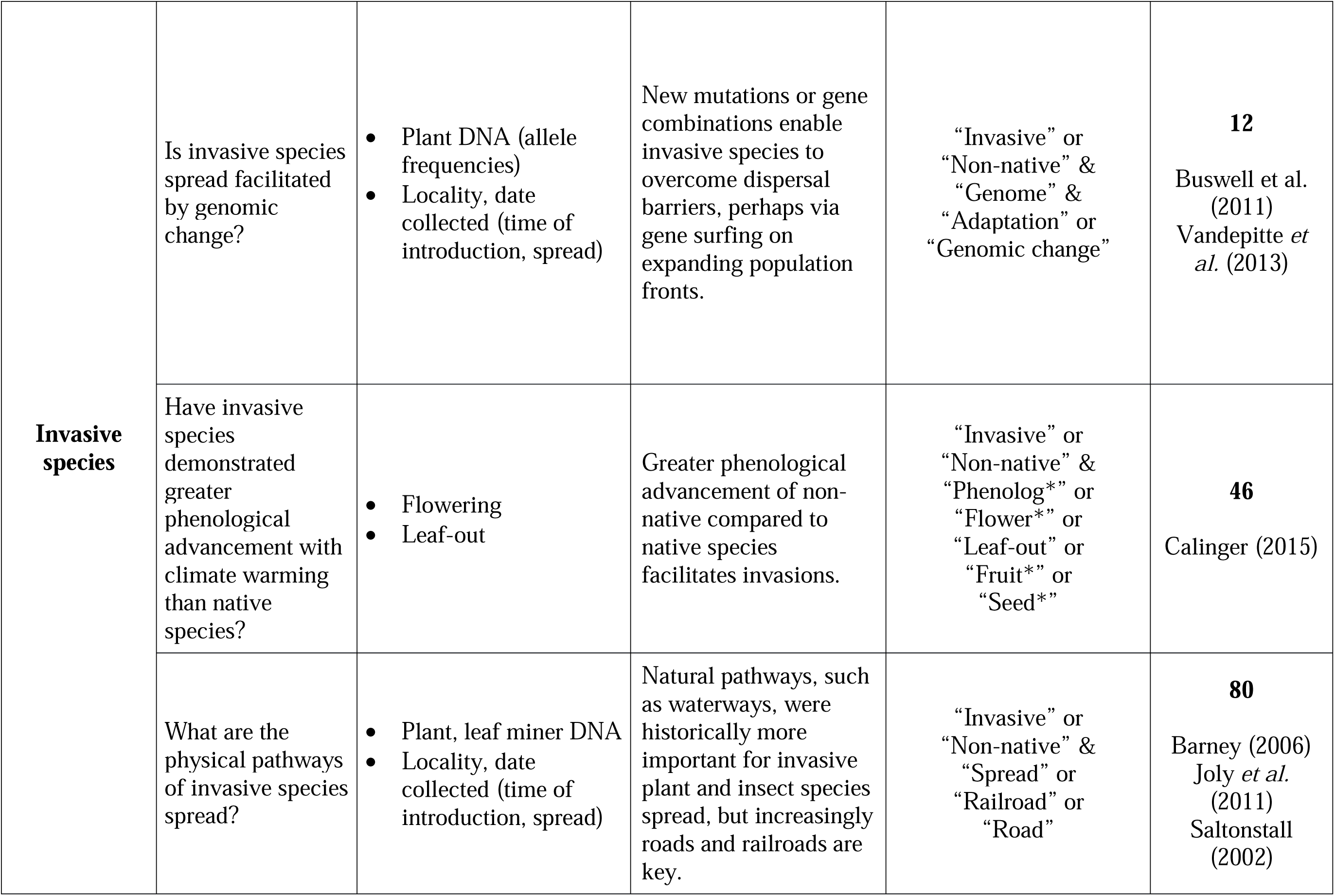
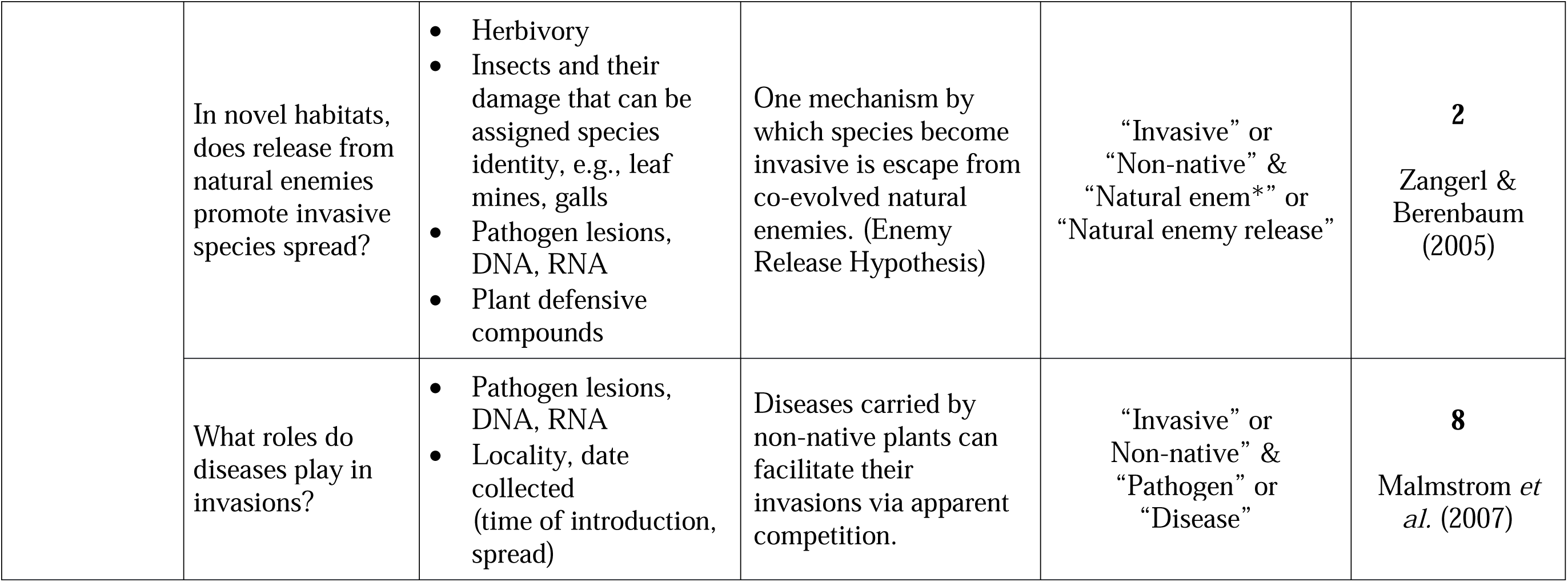
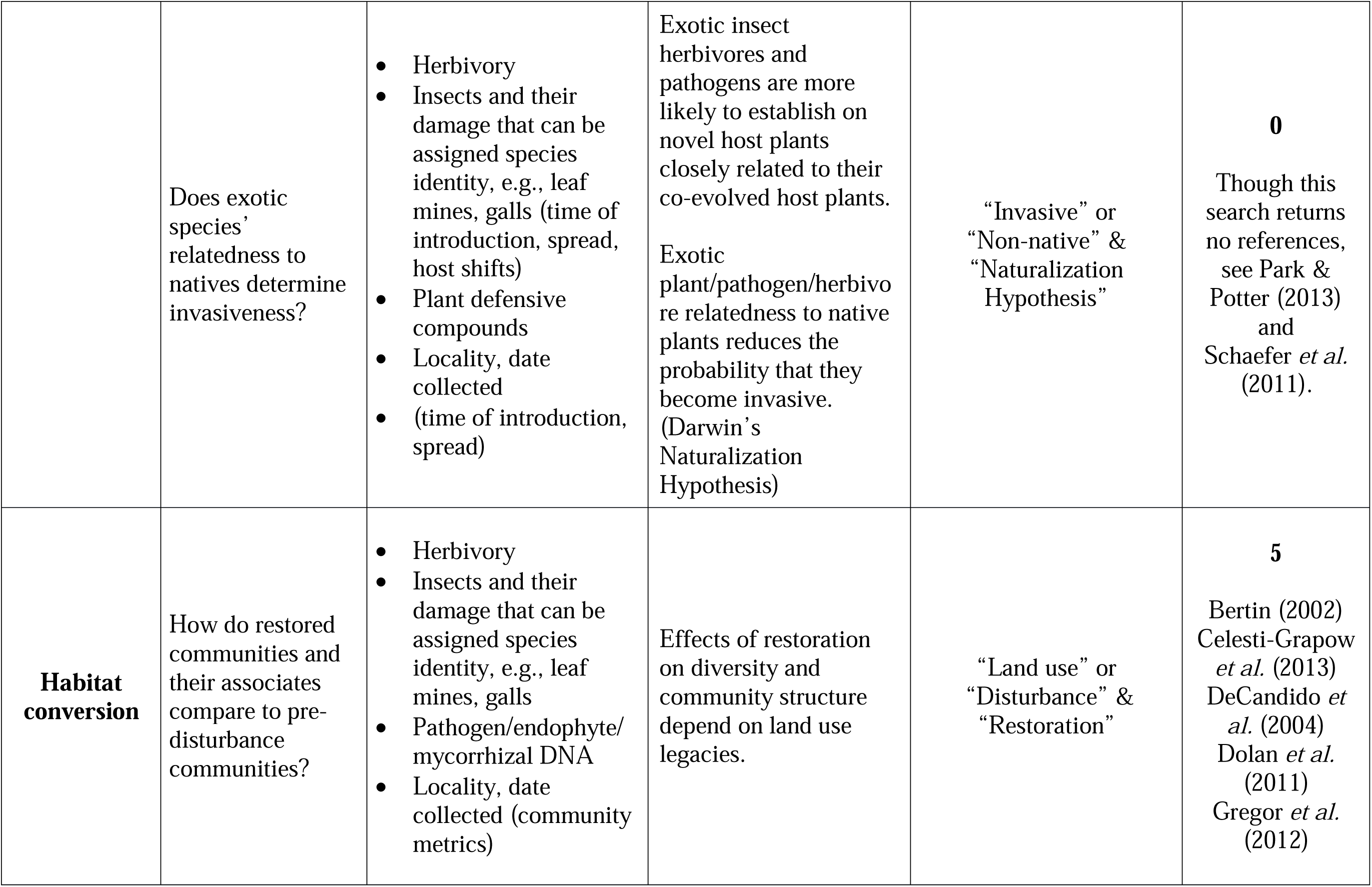
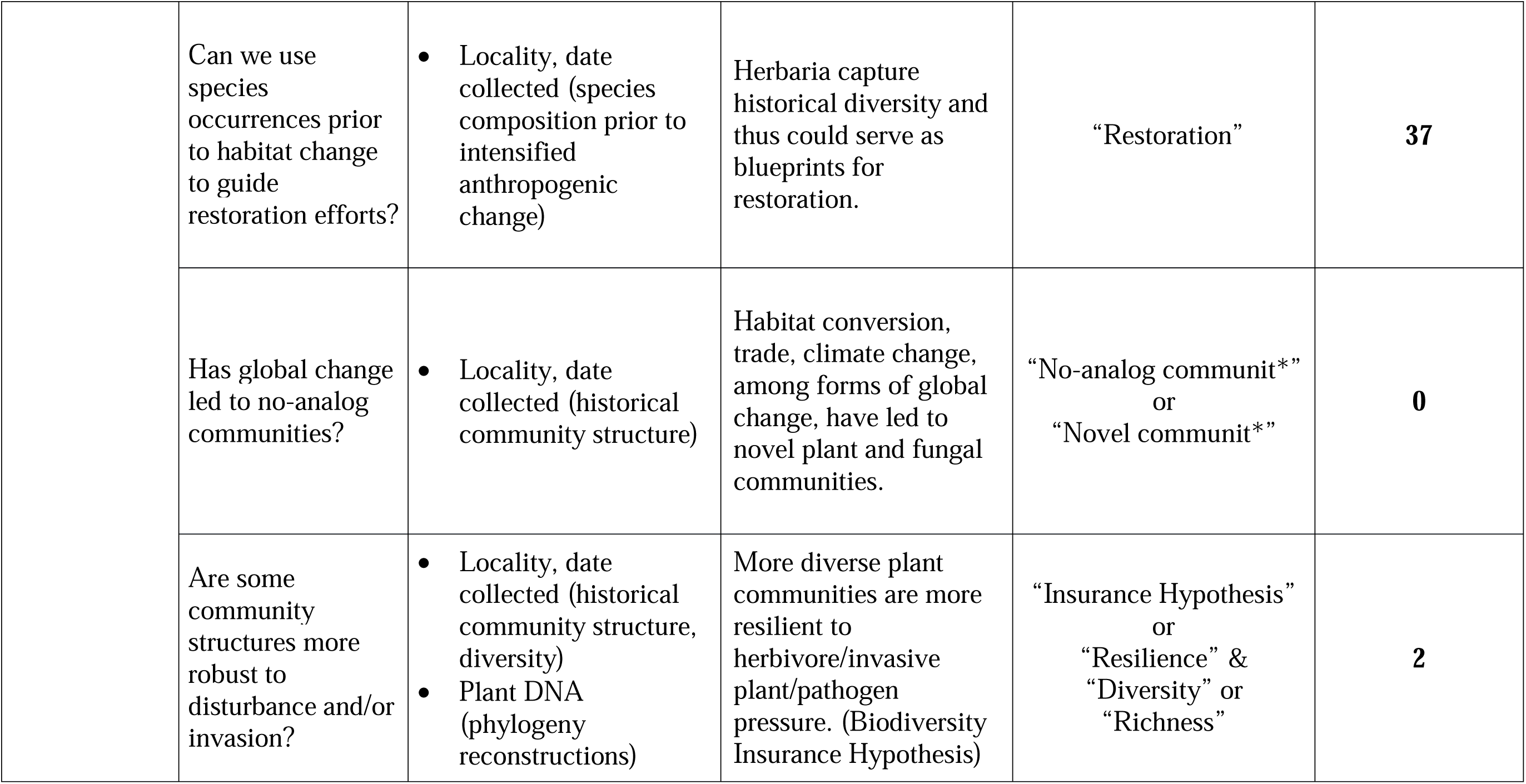
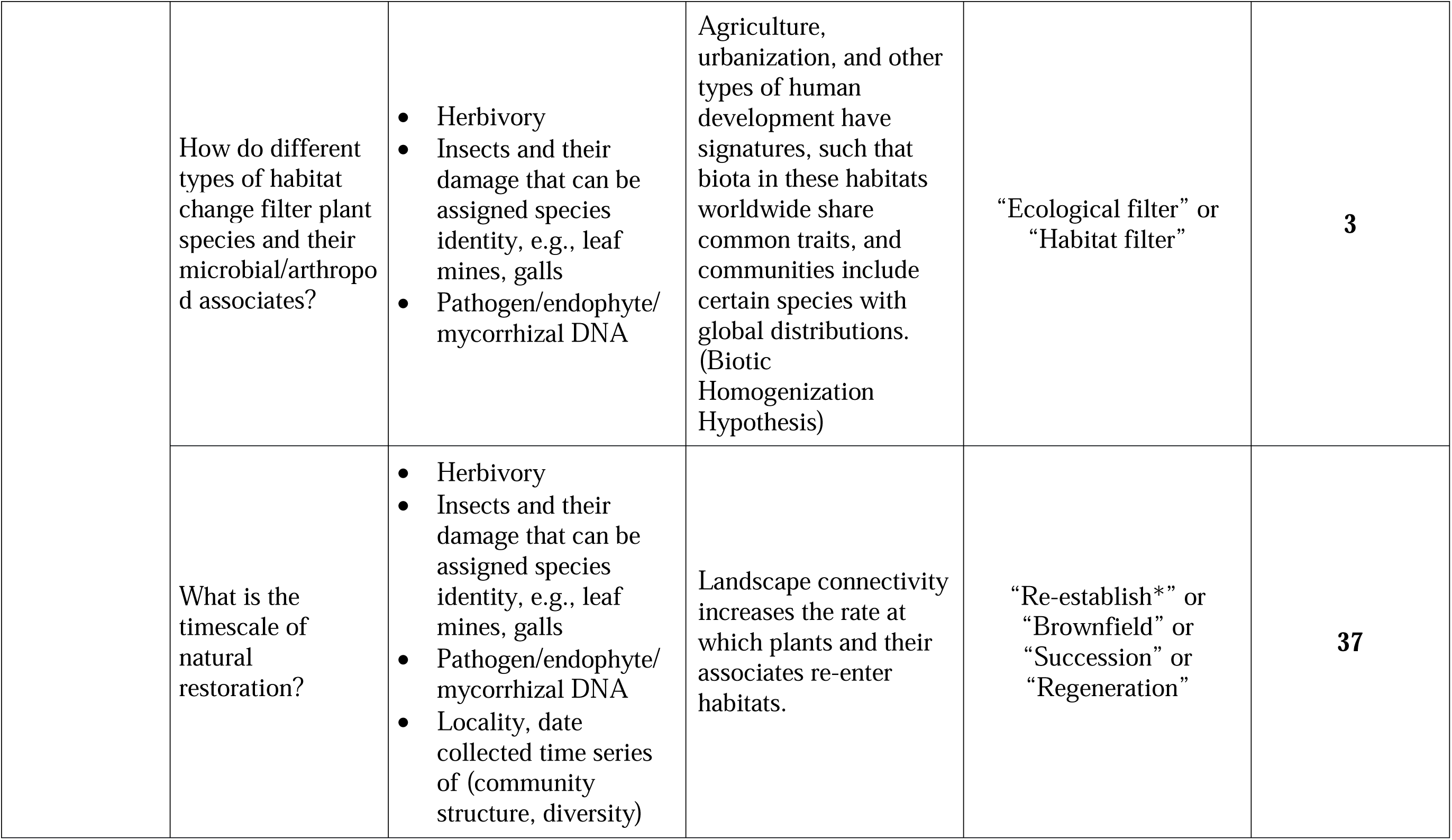
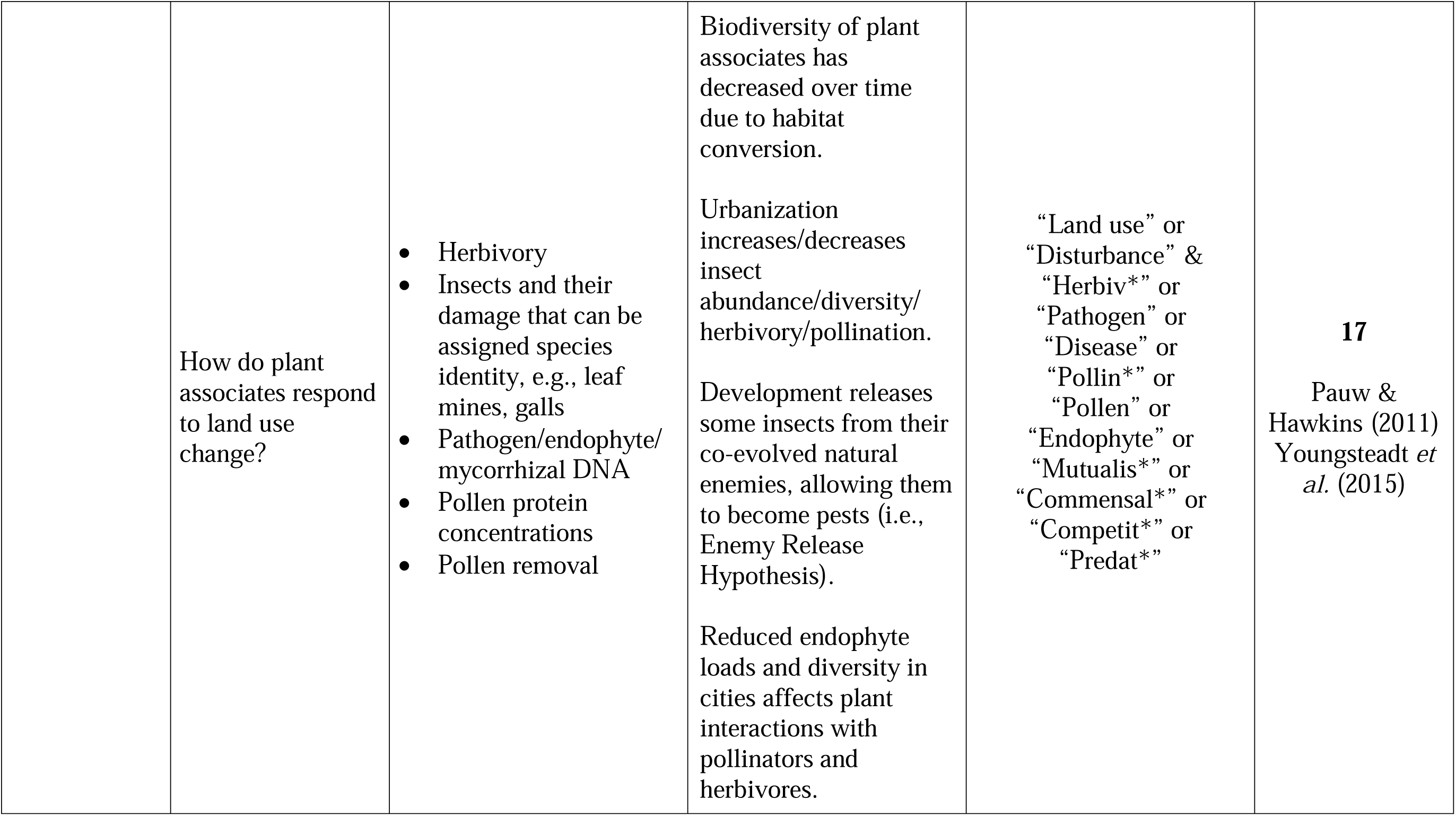
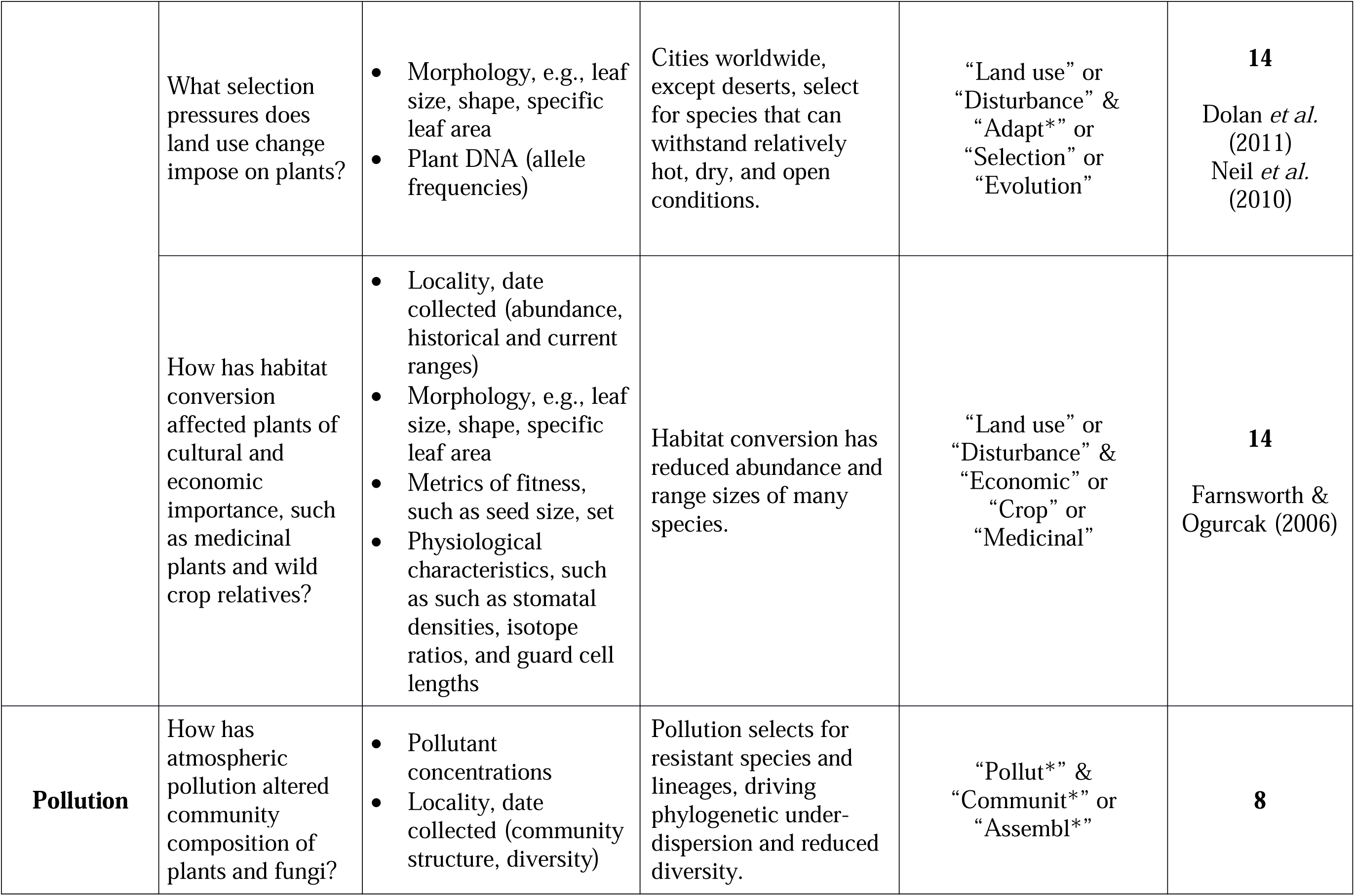
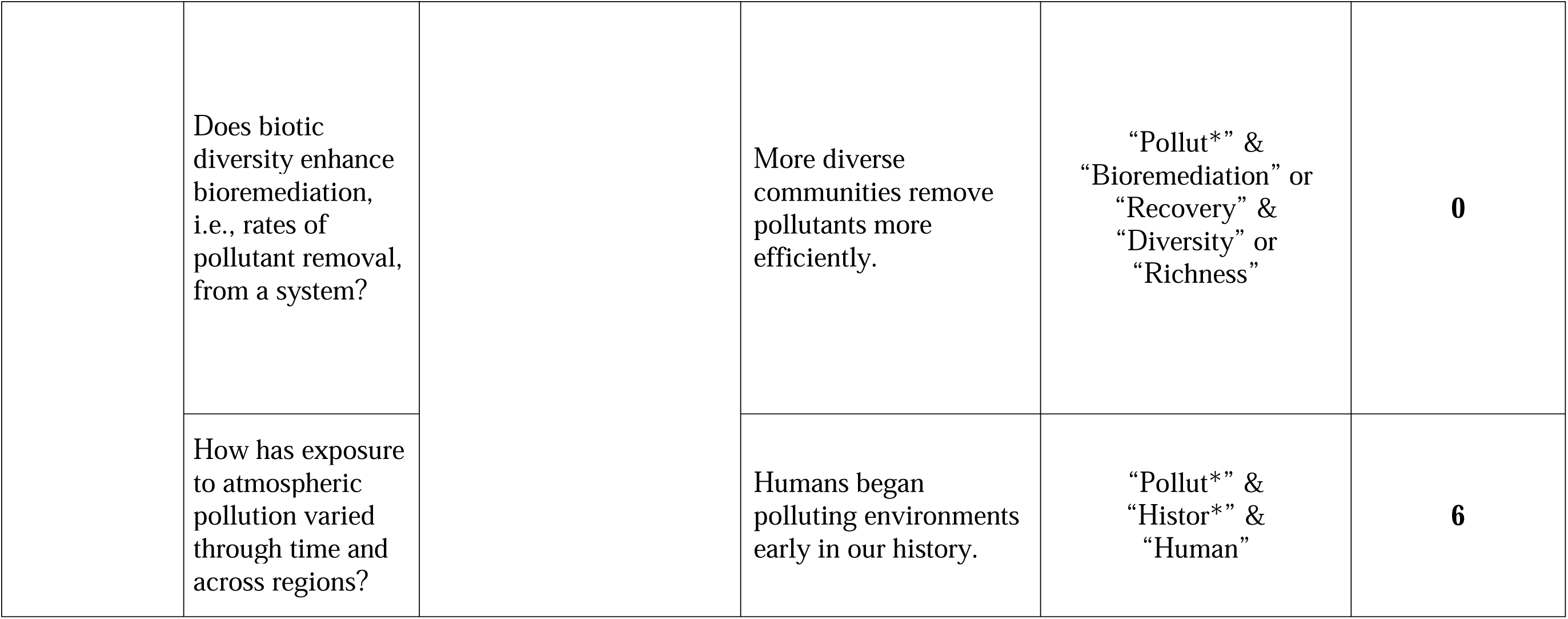
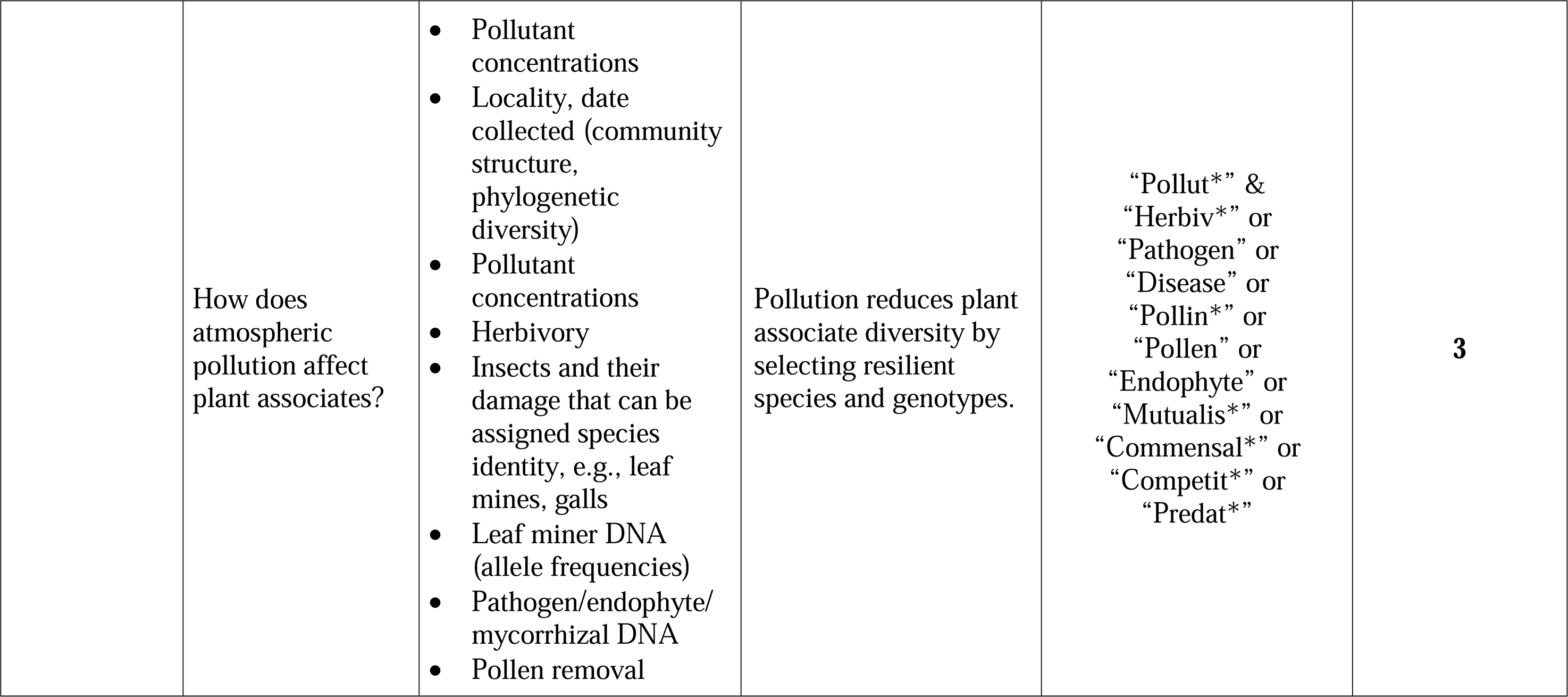
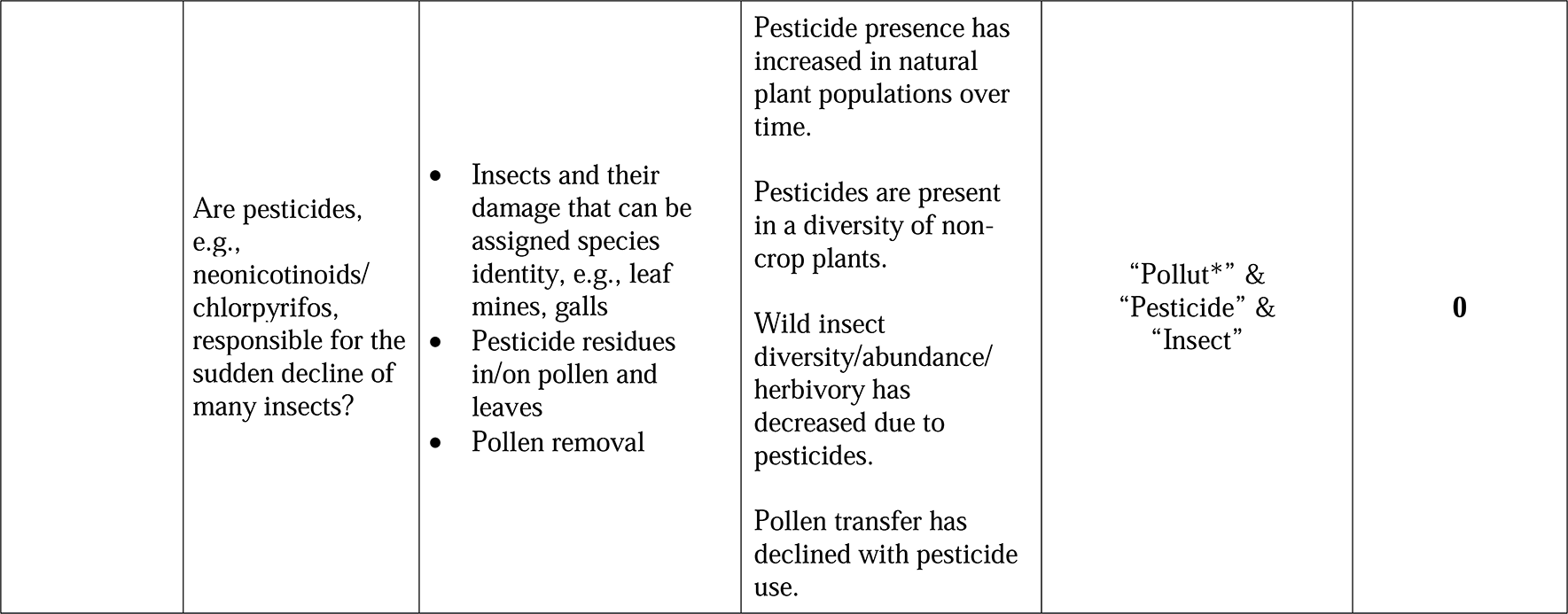
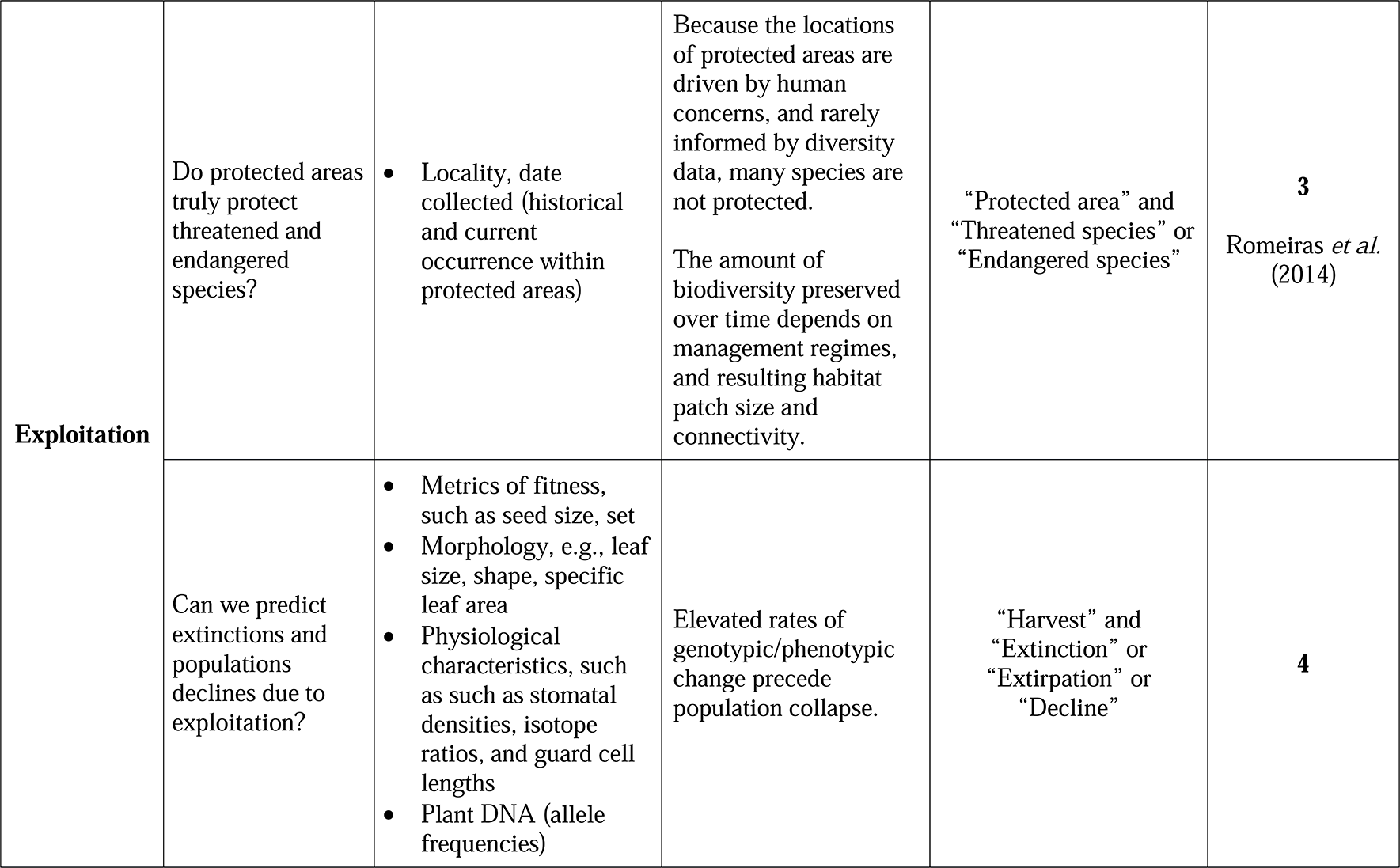
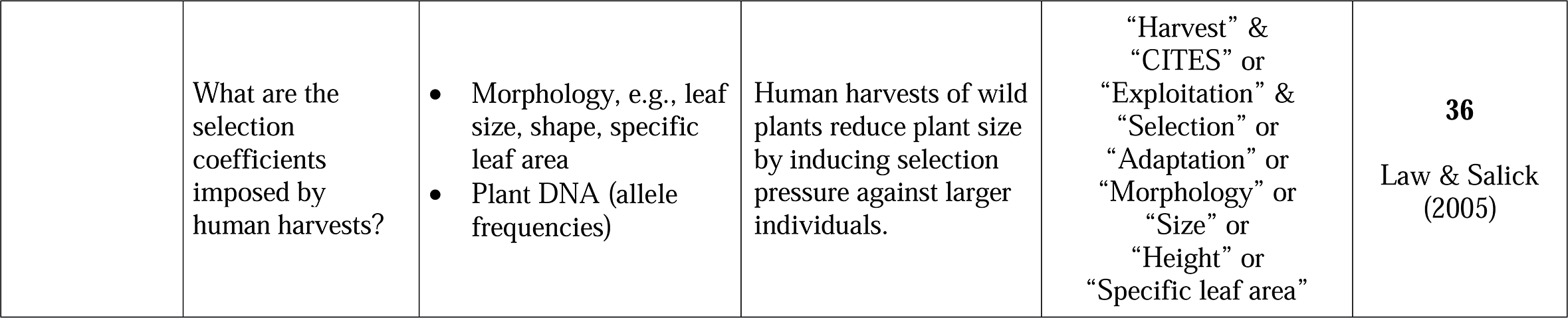
Key research questions and hypotheses from the global change literature for which herbaria may be well-suited as a data source. Most of the research questions apply to multiple global change drivers and are grouped by the driver that has received the most interest. These suggestions are biased toward plants, for which more diverse herbarium data extraction methods have been developed, though we believe many could also be tested for fungi. We searched Web of Science to estimate the current interest in each research topic. Searches were performed with ‘“Herbari*” &’ the listed keywords. The “Citations” column contains the number of citations retrieved and, when available, iconic, representative examples from the literature.

Given the potential value of herbaria to global change research, it is perhaps surprising that, with the exception of their use as records of species in occurrences and phenology, they have not been more widely used. One explanation is that herbarium data present distinct challenges not present in data gathered from experiments and field observations. Some of these are associated with the vast and dispersed nature of herbarium collections. For example, selecting appropriate focal taxa is critical in any ecological or evolutionary study. While collections contain millions of specimens, they often lack easily accessible digital records. This makes determining which taxa are well-represented within and across collections difficult. Other challenges arise as a result of biases; plants are more frequently collected at certain times of year to capture flowering or fruiting, and collection effort has been uneven across space and time (Meyer *et al.*, 2016; Daru *et al*., *In Press*). Challenges also arise because of preservation artefacts; DNA degrades, specimens lose coloration over time, and insects often eat pressed plants housed within museums. Extracting credible data from collections presents significant and complex challenges, and, thus, much of their potential has remained untapped (Davis *et al.*, 2015; Meyer *et al.*, 2016).

In contrast to previous perspectives, which have highlighted the potential of herbaria to inform our understanding of species phenology and geographic distributions, particularly with regard to climate change (Lavoie & Lachance 2006; Primack *et al.*, 2004; Pyke & Ehrlich 2010; Johnson *et al.*, 2011; Lavoie 2013; Vellend *et al.*, 2013; Willis *et al.*, 2017a), here, we explore the broader applications of herbarium specimens to global change research. First, we consider less common applications to global change hypotheses, emphasizing novel methods. Next, we discuss current limitations of using herbarium collections for these purposes by identifying research gaps and challenges associated with collecting and analyzing herbarium data. To illustrate how some such challenges may be overcome, we present a case study focused on species interactions, an aspect of global change biology that is data-poor, but for which herbarium data may be suited despite biases in the underlying collections. Finally, we outline novel future research directions and suggest curation priorities with the goal of positioning herbaria as primary data repositories for ecological and evolutionary research on the effects of global change.

## CONTRIBUTIONS OF HERBARIA TO GLOBAL CHANGE RESEARCH

Interest in applying herbarium data for investigating global change has grown in recent decades. While climate change has been the focus of most studies, use of herbarium data is slowly permeating other areas of global change biology research. Here, we briefly review the ways in which these data have been employed to examine biological responses to the five key drivers of global change and biodiversity loss: climate change, habitat conversion and degradation, invasive species, and—where studies are available—pollution, and overexploitation. We categorize research addressing these drivers into three types of biotic responses: shifts in distributions and population sizes, changes in physiology or morphology, and altered ecological interactions. Within these categories, we consider each of the five key drivers of change ordered by how thoroughly they have been investigated. We highlight particular studies that we consider to be most innovative or that best represent a general approach and apologize in advance for not being able to cite all the many excellent publications that have used herbarium data and which have addressed a wide range of important and exciting questions.

### Shifts in distributions and population sizes

Herbarium specimens typically include collection localities and therefore serve as occurrence records that are now widely used to parameterize species distribution models and to understand the effects of recent global change on species distributions (Feeley 2012; Calinger 2015; D’Andrea *et al.*, 2009). For many species, herbarium specimens have revealed plant species range-shifts both upwards in elevation and pole-wards in latitude in response to recent warming (Feeley *et al.*, 2013), with some species’ ranges contracting while others expand (Feeley 2012). The use of herbarium specimens in this context is now widespread. Though most studies have focused on terrestrial vascular plants, some have included algae (Riera *et al.*, 2015) and species from aquatic systems (Wernberg *et al.*, 2011; Yaakub *et al.*, 2014), highlighting the taxonomic and functional diversity represented in herbaria.

Herbaria occurrence records have also revealed the effects of habitat conversion on species composition, ranges, and abundance. In developed countries, this research has focused on urbanization, arguably the most profound form of global change in these regions (United Nations 2007). Herbarium specimens are among the only floristic records for these areas prior to development and have revealed that urbanization drives native species declines across many cities (e.g., Bertin 2002; DeCandido *et al.*, 2004; Dolan *et al.*, 2011; Gregor *et al.*, 2012; Celesti-Grapow *et al.*, 2013). Herbaria can also provide insights into urban filters on plant communities and have demonstrated that urbanization reduces the occurrence of species associated with wetland habitats (Bertin 2002; DeCandido *et al.*, 2004; Dolan *et al.*, 2011; Gregor *et al.*, 2012; Celesti-Grapow *et al.*, 2013). Unfortunately, while habitat loss through urbanization and deforestation in remote, highly biodiverse regions, such as the wet tropics, is likely a major driver of extinction (e.g., see Wearn *et al.*, 2012), herbarium specimens from many of these regions are relatively sparse (Meyer *et al.*, 2016) and thus provide poor baselines from which to derive estimates of biodiversity change (Feeley & Silman 2011). Nonetheless, herbarium data can help identify species in decline and regions that may provide refuges (Farnsworth & Ogurcak 2006; Romeiras *et al.*, 2014), as well as capture the interactions between climate change and habitat conversion, notably to demonstrate how human land use limits the extent to which species can track their climatic niches (Feeley & Silman 2010).

While many native species are declining, non-native species are increasing in abundance, and a small proportion of these have become invasive (Mack *et al.*, 2000). Herbaria provide historical records of invasive species spread through time and across space (Lavoie *et al.*, 2007; Crawford & Hoagland 2009) and thus can help identify the mechanisms by which non-native species have dispersed from continent-to-continent and expanded their geographic ranges. Analyses of plant occurrence records from herbaria have revealed that human transportation networks, notably paved roads and railroads, are important pathways for invasion (e.g., Barney 2006; Joly *et al.*, 2011). With recent advances in molecular techniques, it is now possible to map the spread not just of species but also particular genotypes using herbarium data, as has been illustrated for the common reed, *Phragmites australis* (Saltonstall 2002). These fine-scale historical data can provide information on both the rate and direction of spread, revealing likely centers of introduction and regions of high vulnerability.

### Physiological and morphological change

Most species are unable to completely escape global change in space and thus must adapt or acclimate *in situ* or risk extinction. Herbarium specimens can capture physiological and morphological changes reflecting such responses, including shifts in morphology (Law & Salick 2005; Leger 2013), timing of life histories (Kharouba & Vellend 2015), and physiology (Miller-Rushing *et al.*, 2009). In a small but growing number of studies, physiological and morphological data, such as on plant leaf size (Guerin *et al.*, 2012), stomatal densities (Miller-Rushing *et al.*, 2009), carbon and oxygen isotope measurements (Miller-Rushing *et al.*, 2009; Bonal *et al.*, 2011), and specific leaf area (Reef & Lovelock 2014), have been extracted and provide insights into changing photosynthetic rates and leaf palatability for herbivores, for example.

The most studied of these shifts with herbarium data are changes in plant phenology. Flower counts from herbarium specimens have revealed advances in peak flowering of approximately 2.4 days for each degree Celsius rise in temperatures (Calinger *et al.*, 2013), and similar advances due to the urban heat island effect (Primack *et al.*, 2004) that are particularly pronounced in ephemeral species (Neil *et al.*, 2010). Researchers have also turned to herbaria to identify potential cues driving plant phenology, revealing interactive effects of temperature, precipitation, and latitude on the timing of flowering (Matthews & Mazer 2016). Such data have proven extremely valuable as both a biotic index of climate change, and as a record of biotic responses to climate warming. The extraction and application of phenological data from herbaria has been reviewed extensively elsewhere (Miller-Rushing et al. 2006; Willis *et al.*, 2017a).

Plant phenology is just one response to changes in atmospheric chemistry. Concentrations of pollutants in the atmosphere, including heavy metals, anthropogenic nitrogen, carbon dioxide (CO2), and other greenhouse gases have varied over time, with increases surprisingly early in modern human history (Renberg *et al.*, 1994; Steffen *et al.*, 2007). However, because historical data are sparse, and pollutant sources are often diffused, changes in pollution can be hard to track using traditional ecological approaches. Herbarium specimens serve as records of pollutant variation over space and time, helping link species’ exposure to their responses. For example, epiphytes, which accumulate atmospheric nitrogen, but do not uptake nitrogen from soil, can serve as bioindicators of nitrogen pollution (Stewart *et al.*, 2002). Mosses similarly serve as bioindicators of atmospheric metals (Weiss *et al.*, 1999), and lichens serve as bioindicators of various pollutants, including a diversity of oxidants (Sigal & Nash 1983).

Data from herbaria can also be used to identify the traits associated with invasiveness. A key question in invasion biology is why few species become invasive while most non-native species remain at low abundances (Sakai *et al.*, 2001). By capturing physiological and morphological data, herbarium studies have shown that adaptation to local conditions can facilitate invasiveness (Vandepitte *et al.*, 2014) and, consistent with other types of data (Wolkovich & Cleland 2011), that some invasive species are able to better track temperature and thus take advantage of earlier springs resulting from warming (Calinger 2015).

### Shifts in Ecological Interactions

Ecological interactions are increasingly recognized as moderating species responses to global change (Gilman *et al.*, 2010; Zarnetske *et al.*, 2012), but empirical data on this topic are sparse. Herbarium specimens have served as records of the interactions between plants and their associates, revealing how these interactions have shifted over time. These plant associates, including insect pollinators, herbivores and pathogens, are sometimes preserved on leaves (Lees *et al.*, 2011) and branches (Youngsteadt *et al.*, 2015) or are preserved as DNA or RNA (Malmstrom *et al.*, 2007). In addition, flowers and leaves of herbarium specimens can contain signatures of interactions, such as the accumulation of defensive compounds induced by insect herbivores (Zangerl & Berenbaum 2005) and pollen (Ziska *et al.*, 2016).

The trace-record of interactions between plants and pollinators, herbivores, and pathogens captured on herbarium specimens could be used to address a topic that has attracted much interest in climate change research: potential phenological asynchronies and ecological mismatches between associated species resulting from differential responses to warming climates (Post *et al.*, 2008; Both *et al.*, 2009). Phenological asynchronies occur if the direction, rate, or magnitude of change differs between associated species, if phenological responses differ in space, or if responses within a single trophic level are varied, such that phenological tracking is not possible (as discussed in detail elsewhere, e.g., Hegland *et al.*, 2009). Occurrence data from herbaria and contemporaneous insect collections have suggested that climate change might lead to asynchronies between some butterflies and their adult food plants (Kharouba & Vellend 2015). Other types of ecological mismatches can also disrupt mutualisms. Corolla tube length data from herbarium specimens, along with corresponding bumble bee collections that provided data on bee tongue length, showed that bees and flowers may be ecologically mismatched in Colorado, USA; bee tongues lengths have decreased over time, while corolla tube lengths have remained constant (Miller-Struttmann *et al.*, 2015). However, there are few studies along these lines, and we expand on the potential of herbarium data for studying phenological asynchronies in the following section.

Similar data can serve as records of changing species interactions due to habitat conversion, pollution, and invasions. Anthropogenic disturbance of a habitat can expose species to novel conditions that disrupt co-evolved interactions. By rehydrating orchid flowers from herbaria and counting the pollinaria (pollen clumps and associated tissues removed by bees), Pauw & Hawkins (2011) demonstrated that the local decline of an orchid during urbanization was driven by reduced pollination. Youngsteadt *et al.* (2015) counted scale insects on stems of herbarium specimens and demonstrated that the urban heat island effect and natural warming cycles in forests are associated with elevated abundance of a scale insect herbivore, suggesting that ecological responses to warming in cities could predict the effects of climate change on insect pests. In one of the few herbarium studies to explore how pollution affects species interactions, Ziska *et al.* (2016) analyzed pollen preserved in herbarium specimens and demonstrated that elevated CO2 concentrations reduced pollen protein concentrations, altering nutrient availability for pollinators, with possible fitness consequences for both pollinators and plants. In a separate study, Zangerl & Berenbaum (2005) provided support for the natural enemy release hypothesis of plant invasion (Keane & Crawley 2002; Mitchell & Power 2003) by measuring exotic plant defensive compounds before and after introduction of their co-evolved herbivores. They found evidence that invasive plants can escape their insect herbivores—and the need to manufacture expensive defensive compounds in response to them—when first introduced into new habitats.

## GAPS IN THE APPLICATION OF HERBARIUM DATA

### Overview

We have provided a brief overview of the diverse applications of herbarium data for understanding biotic responses to global change. However, many of these data have only recently become commonplace in ecological studies and we suggest their potential has not been fully realized. Here, we identify significant gaps in current uses of herbarium specimens in global change biology (Table 1). As we indicate above, data from herbaria have been used extensively to explore plant distributional and phenological shifts in response to climate change. However, herbarium specimens have been used less frequently to study plant responses to three of the five key drivers of biodiversity loss: habitat conversion, pollution, and overexploitation. By many estimates, habitat conversion currently drives most terrestrial species extinctions (Millennium Ecosystem Assessment 2005), making this a particularly notable omission. With these overarching themes in mind, we suggest future research directions we believe could benefit from using herbarium data.

### Guidance for ecosystem management and restoration

Herbarium specimens are rare records of historical biodiversity and thus could help guide ecological restoration. A few studies have highlighted this potential, mostly within the context of urban environments (DeCandido *et al.*, 2004; Atha *et al.*, 2016). However, less common are examples of using herbarium data to explore other types of habitat conversion, such as agricultural expansion and deforestation, much less integrate such findings into restoration programs. As is the case for urbanization, these types of habitat transformations are likely to have nuanced effects on local biodiversity that are not well understood but which may be captured by collections. Herbarium specimens can additionally be used to determine areas and species of conservation priority, as has been demonstrated for timber species in Angola (Romeiras *et al.*, 2014). We suggest there may be many opportunities along these lines for conservation of plants of economic and/or cultural importance, including both timber species and wild crop relatives, which are often well-represented in herbaria, with many specimens collected prior to intensification of global change.

### Markers of plant physiological change

Specimens within herbaria provide more than just records of occurrence across space and time; they also bear the imprint of past environments, including information on genotypic and physiological shifts. For example, herbarium specimens could be employed to collect long-term data on shifts in traits correlated with photosynthetic rates and nutrient concentrations across taxa and habitats. These are ecosystem variables for which we have little historical data, but which have important consequences. For example, leaf gas exchange rates can affect carbon balance (Bonan 2008), and the net effects of global change, particularly climate change, on this and similar ecosystem processes is a topic of great interest (Clark 2004; Wu *et al.*, 2011). Increasing CO2 levels are expected to increase photosynthetic rates in plants, increasing net carbon storage in forests (Ainsworth & Long 2005). However, long-term experiments to evaluate the relationship between CO2 and photosynthetic rates have been in place for less than 20 years (Norby *et al.*, 2016) and do not incorporate other recent global changes that may have interactive effects on photosynthesis, most notably warming, changes in soil moisture, and nitrogen deposition. While methods have been developed to extract such data from herbarium specimens—for example, quantifying proxies for gas exchange preserved in leaves of pressed plants (Miller-Rushing *et al.*, 2009; Bonal *et al.*, 2011)—and are not new (Woodward et al. 1987), they have much unrealized potential in global change research.

### Records of changing species interactions

We suggest that herbarium specimens could provide unparalleled insights for understanding shifting species interactions in the Anthropocene. Above, we outlined a few studies along these lines, but we believe the data within herbarium specimens has not been fully exploited. For example, presently, there is little consensus on whether climate change induced phenological asynchronies are common or rare, in large part because data are sparse. It is, however, possible to pair herbarium records with collections of species with which they are tightly associated, such as their pollinators. This approach is potentially powerful, but requires substantial data. It is possible, therefore, that for many species, collections will not yield data of sufficient spatial and temporal resolution to quantify phenology for associated species. An alternative approach is to draw indirect inferences on species interactions using data from only one partner; observations of pollen, for example, can provide information on plant-pollinator interactions (Pauw & Hawkins 2011; Ziska *et al.*, 2016). Such methods can be more easily expanded to systems for which corresponding collections on interactors are sparse or unavailable.

In a number of studies, researchers have used similar approaches to explore effects of global change on antagonistic interactions between plants and their associates by extracting data on herbivores, herbivory, and plant pathogens (Malmstrom *et al.*, 2007; Lees *et al.*, 2011; Youngsteadt *et al.*, 2015; Syfert *et al*., 2017). However, methods remain relatively underdeveloped. Plant-insect interactions might be of particular interest, because insects are ectothermic, and, thus, their abundances are expected to shift with global climate warming in many cases (Kingsolver *et al.*, 2013). Herbivory is also ubiquitous among present day plants (Turcotte *et al.*, 2014), has driven evolution of much of the biodiversity on earth (Futuyma & Agrawal 2009), and has significant economic consequences in agriculture and forestry (Oerke & Dehne 2004). Despite its importance as an ecological process, we are aware of few studies quantifying plant-insect interactions on individual specimens (but see Morrow & Fox 1989).

## HERBARIA AS NOVEL DATA SOURCES: LIMITATIONS AND CHALLENGES

We have shown that herbarium data can be applied to diverse topics and have suggested gaps that warrant future exploration. Here, we outline challenges these data present and, when available, approaches to reduce obstacles to their use, which we believe is a key hurdle to using herbarium data in global change research. We consider some general methods that could be applied widely to herbarium and biological collections data. Finally, we provide a table detailing potential challenges posed by specific types of herbarium data, along with suggested ways to approach these challenges (Table 2).

**Table 2.** Herbarium data for global change research. Here, we highlight the different data types, challenges to their use, and potential solutions to overcome these challenges. We focus here on the data types and uses we have discussed in the main text; other applications may be subject to additional challenges (and potential solutions).

| Herbarium data | Challenges | Potential solutions |
| --- | --- | --- |
| All data | Biases over space, time, and phylogeny | Careful focal herbaria/species selection aided by digitized specimens<br>Subsampling and techniques for inference with biased data, and statistical methods such as machine learning |
| Occurrence (locality, date collected) | Lack of recorded absences | Statistical tools for simulating or otherwise analysing presence-only data, which are already well-developed for species distribution models and may be employed for other areas of research. |
|  | Coarse-level geographical data. | Careful focal herbaria/species selection aided by digitized specimens<br>In some cases, it may be possible to address hypotheses with environmental data at coarse scales, e.g., temperature data averaged at the county |
|  |  | level in the U.S. |
| Phenology | Specimens are most likely to be collected at peak flowering times, thus missing early season phenological events and times of first event. | • Use statistical estimators to infer timing of first events from a sampled distribution (Pearse *et al.* In Press). |
|  | Collectors tend to retrieve specimens from near roadsides (Daru *et al.* In Press), which could affect phenology. For example, life events may be advanced relative to plants in more natural areas due to the urban heat island effect or delayed due to drought-like conditions. | • When comparing across space, time, and/or phylogeny, relative measures may be sufficient to address hypotheses of interest.<br>• For studies that do require absolute measurements, use specimens with geolocations and model potential biases (e.g., urbanization).<br>• Additional studies are needed to assess the effects of local collection biases on global change data, especially for plant traits that are sensitive to temperature, such as phenology. |
|  | For leaf-out, categorical criteria, such as pubescence, can be used to determine when specimens have newly flushed leaves (see Everill *et* | • Continuous criteria can be developed, such as leaf size relative to fully flushed leaf size (also see Everill *et al.* |
|  | *al.* (2014)). However, many species do not have unique qualities associated with new leaves. | 2014), but such criteria introduce considerably more work than do categorical characteristics. |
| Other traits (morphology, physiology, fitness, and microbiomes) | Botanists may be more likely to collect specimens from more vigorous individuals or branches with greater seed set, seed size, leaf size, water use efficiency, etc. | • When comparing across space, time, and/or phylogeny, relative measures may be sufficient to address hypotheses of interest.<br>• Cross-validate recent collections with observational data to ascertain, and thus allow correction for, potential sampling biases. |
|  | Key tissues, morphological structures, or developmental stages may not be sampled or may be damaged. For example, roots are necessary for mycorrhizal studies but are not present on all specimens. | • Check digital images and select intact specimens.<br>• If alleles for a trait are known, it may be possible to genotype a sample without a visible phenotype and thus avoid the need for the actual structure possessing the trait of interest.<br>• For certain species, roots can be easily collected and are thus present on many herbarium specimens. Overcoming this challenge should be a matter of |
|  |  | choosing appropriate focal taxa. |
| Pollination | Matching data on pollinators are often not available. | Score pollination on herbarium specimens. This will only be possible for groups that have distinct pollen-holding structures (pollinaria), such as many orchids and milkweeds. |
|  | To build matching plant-pollinator collections, one needs to know which pollinators are associated with which plants. | Identify plants via pollen morphology or DNA preserved on pollinator specimens. |
| Antagonistic interactions (herbivory, disease) | Botanists may be likely to collect less damaged specimens. | When comparing across space, time, and/or phylogeny, relative measures may be sufficient to address hypotheses of interest.<br>Cross-validate recent collections with observational data to ascertain, and thus allow correction for, potential sampling biases. |
|  | Difficulty differentiating between taxa. For example, it may be difficult to identify taxa that created galls and leaf mines because of specimen degradation. | Combined morphological and molecular methods may improve identifications.<br>Focus on species with distinct, |
|  |  | well-preserved gall, mine, and pathogen damage morphologies.<br>• Direct sequencing of associates if preserved on plant specimens, for example, using DNA barcodes for species identification. |
|  | DNA, RNA degradation | • For most species, RNA will be too degraded. However, Malmstrom *et al.* (2007) have shown it is possible to extract usable RNA in some (perhaps rare) cases.<br>• Next generation sequencing techniques developed for amplifying degraded or ancient DNA<br>• Collection of tissue samples preserved in silica or banked in ultra-low temperature storage. |
|  | Quantifying damage on individual specimens | • We have demonstrated the potential of subsampling specimens to quantify herbivory damage or pathogen lesions. (See the “Limitations and Challenges” section), |
|  |  | analogous to the use of quadrats in vegetation sampling.<br>• In cases where visual signs of herbivory are difficult to quantify, defensive compounds might provide an alternative way to detect damage. |
| Pollution | Choosing appropriate indicator species for pollutants, regions, and time periods of interest | • Indicator species for many pollutants, such as nitrogen and heavy metals, have been established. Digitized records can then help select those with suitable temporal and geographic coverage. |
|  | Pesticide degradation | • Some pesticides degrade faster than others, and recovering pesticide residues may be possible for chemicals that break down more slowly. This area of research has not been explored, and thus methods are not yet developed. |
|  | Botanists may be unlikely to collect in heavily polluted environments or near areas where pesticides have been applied. | • Labels may indicate if specimens were collected near farms, roads, homes, and industry. These metadata can be included in subsequent |
|  |  | analyses. |

### Biases over space, time, and phylogeny

Herbarium specimens are non-randomly collected across space and time, in part because their historical purpose was to document species’ ranges and capture morphological variation within and between species, and not to address ecological questions. Using specimens for global change research requires accounting for this unevenness in sampling over space, time, and taxonomy. This topic has been reviewed recently by Meyer *et al.*, (2016), who analyze biases represented in specimens and observational data, and Daru *et al*. (*In Press*), which includes biases represented in herbarium specimens, specifically. Though the sampling biases described in these publications should serve as a roadmap for those expected more generally in herbarium data, the gaps and biases present across entire herbaria do not necessarily represent those in the subset of data extracted for particular studies. For example, an herbarium may include few species from a particular province in China, but have many specimens for each of these species.

A first step in any ecological analyses is to design appropriate sampling procedures to minimize biases. There is a rich literature on sampling techniques in ecology (see Southwood & Henderson 2009), but these have been largely overlooked when “sampling” herbarium specimens. Instead, when deriving data from specimens in ecological research, it is more common to conduct systematic sampling, analyzing specimens within a specified timeframe and/or spatial area rather than considering herbarium specimens as a sample drawn from a larger population. Now that centralized databases contain millions of specimens, traditional ecological tools, such as stratified random resampling or rarefaction, may allow researchers to minimize biases. In addition, researchers can focus analyses on the richer parts of collections, thus reducing noise and bias introduced by taxa, time periods, or locations for which few specimens are available. For example, if we were to sample a particular plant species across the northeastern US (Fig. 1b; the state in the bottom right is Massachusetts, and the state below it and to the left is Connecticut), we may find that more recent samples are available for Connecticut than for Massachusetts, a known pattern given the historical idiosyncrasies of institutions and collections in the region. Therefore, if we were interested in testing a global change hypothesis that requires a time series, we may decide to focus our sampling in Connecticut to take advantage of its time series and avoid the uneven temporal coverage across space in Massachusetts.

Even when sampling procedures are designed carefully, however, it may not be possible to select specimens evenly across all axes of variation. Numerous modern statistical tools are available to account for uneven sampling and non-independence of data. These include, notably, spatial regression techniques, such as spatial autoregressive models, spatial lag models, and spatial error models (Plant 2012), that account for spatial non-independence in data. Various phylogenetic comparative methods, including independent contrasts (Felsenstein 1985) and phylogenetic generalized least squares regression (PGLS), are available for incorporating phylogenetic non-independence among taxa. There is also a large literature on null models in ecology (Gotelli & Graves 1996) that details methods for subsampling data to incorporate potential biases into null models, thus holding bias constant when testing for relationships among variables of interest. New machine learning tools and Bayesian techniques (notably in programs such as Stan, WinBUGS, and JAGS) allow for the analyses of complex, hierarchically structured, and incomplete datasets and are suited to analyzing large, sparsely sampled data, all common features of collections data. Many of these methods are already well developed for modelling species distributions—e.g., Maximum Entropy, Generalized Linear and Additive Models, Boosted Regression Trees, and Random Forest (Elith & Leathwick 2009)— though they are not yet integrated into ecology more generally (Thesen 2016).

### Data extraction and validation

Once appropriate herbarium specimens are selected, depending on the analysis, it may then be necessary to design sampling strategies to extract data from within individual herbarium sheets. Like any other unit, such as a plot, agricultural field, or transect, individual herbarium specimens can be subsampled to quantify features that may be hard to measure across entire specimens, such as data on arthropod and pathogen associations, or other micro-traits including stomatal density, cell shape etc. In some cases, it may be necessary to measure the surface area of specimens to account for their size, a process that can be automated in programs such as ImageJ ([https://imagej.nih.gov/ij/]), or to estimate specimen density, volume or other dimension to standardize measures for differences in ‘sampling space’.

An important but frequently overlooked next step is to evaluate the reliability of derived statistics. To characterize spatial and temporal biases, researchers have turned to data validation, in which herbarium data are placed on common axes with trusted data. For example, several recent studies have validated the utility of herbarium specimens for phenological research by fitting common models to field observation and herbarium data (Robbirt *et al.*, 2011; Spellman & Mulder 2016). By contrasting flowering data from herbaria to that from observations, Davis *et al.* (2015) showed that herbarium specimens covered more climatic space than observations. Observational data alone might thus provide a more limited estimate of future climate change scenarios than herbarium specimens. This study highlights that traditional data sources also have limitations and gaps in coverage, some of which can be alleviated by including data from herbarium specimens. Similarly, pollution measures derived from herbarium specimens can be calibrated by comparing pollutant concentrations to those from other historical data sources, such as deposits in peat bogs and ice cores (Weiss *et al.*, 1999).

Cross validation with an independent dataset is perhaps the most robust approach for detecting biases in data. However, in many cases, herbarium data cannot be directly compared to independently derived data because no companion data exist, or the collection of such data requires inordinate effort, e.g., the manual cleaning and standardization of multiple datasets. When companion data are unavailable, we suggest another approach for assessing data reliability: comparing summary statistics relevant to the hypothesis to be tested to theoretical expectations from the literature. We demonstrate this approach focusing on insect herbivory, a process that is likely to shift due to global change, but for which few historical data are available (for detailed sampling and statistical methods, see Appendix S1). This is a particularly challenging case study because collectors most likely select specimens that have little damage, so absolute estimates of herbivory are probably strongly biased. Nonetheless, paleontologists have used fossil herbivory, for which data are even more sparse, to assess changes in diversity and abundance of herbivory with climatic changes across epochs (Wilf & Labandeira 1999). Therefore, we suggest that the much higher resolution, more abundant data available from within herbaria should provide at least as much information on contemporary herbivory patterns.

Despite potential biases, as a proof-in-concept example we demonstrate that herbarium specimens harbor diverse types of herbivory damage (Fig. 2). We also provide evidence that they could provide unique longitudinal data on plant-insect interactions (Fig. 3). Herbivory on herbarium specimens has phylogenetic signal (Fig. 3a), with Blomberg’s K value of 0.4 (above zero indicates phylogenetic signal, with a value of one matching to Brownian motion), though our dataset of 20 species precludes formal significance testing. Similarly, the composition of herbivory, i.e., the relative amount of chewing, leaf galls, leaf mines, etc.—is significantly more similar for individuals within than among genera and species (Fig. 3b). These taxonomic and temporal patterns indicate that herbarium specimens may provide adequate data for addressing questions related to how herbivore communities are changing over time and how herbivore pressure varies across plant community members, questions that are of growing interest but for which historical data are limited. In addition, herbivory measured on herbarium specimens is highly skewed, with most specimens showing little or no damage, and a few showing heavy damage (Fig. 3c), as also observed in field data (Turcotte *et al.*, 2014), and accumulates as the growing season progressed (Fig. 3d), which suggests that these data are sensitive enough to capture accumulating herbivory on individual plants through the growing season.

**Fig. 2.**
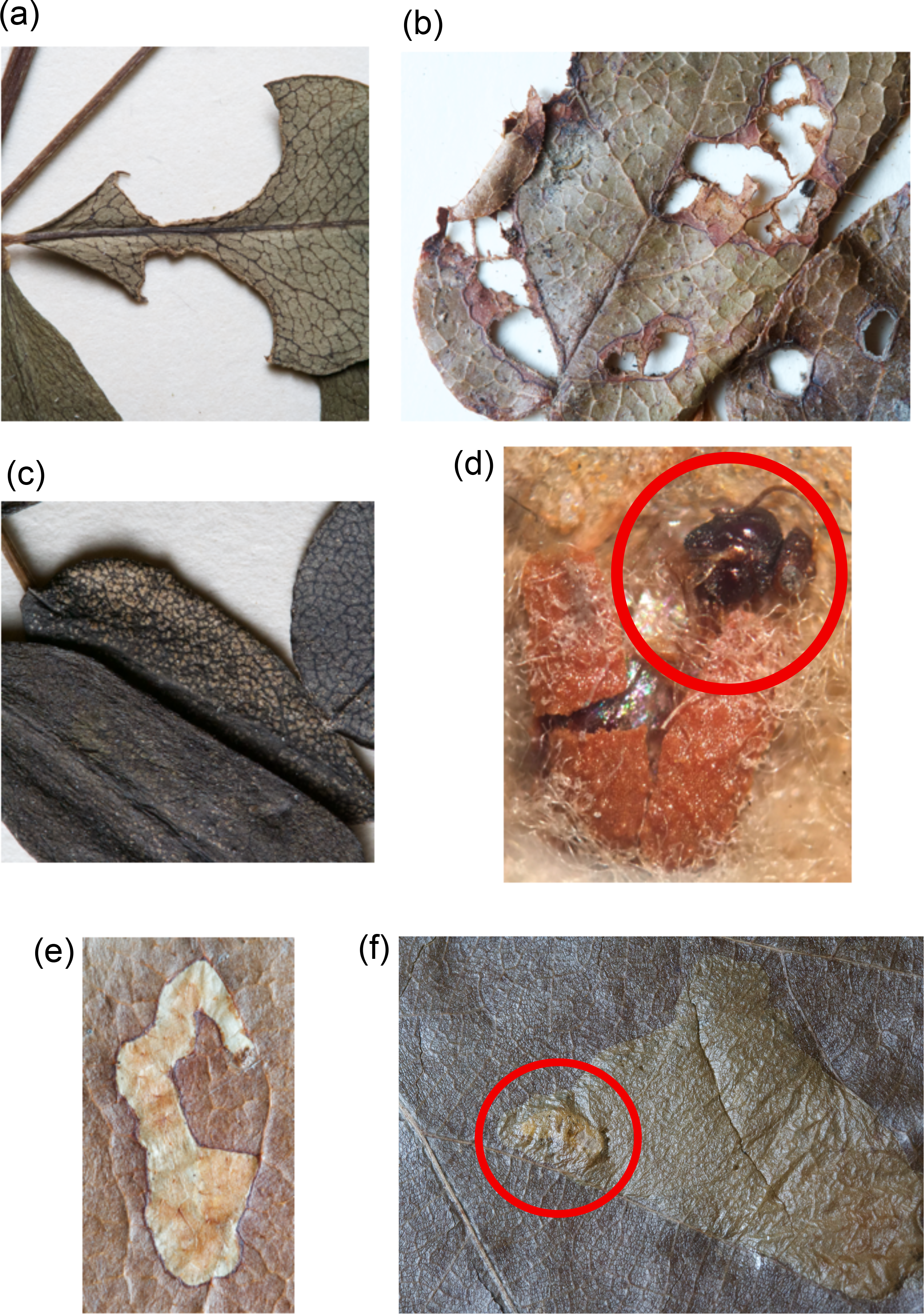
The diversity of insect herbivory preserved on herbarium specimens. We found several types of herbivory on herbarium specimens made by a diversity of arthropods, and quantified herbivory in five categories representing, (a) chewing, (b) skeletonization, (c) stippling, (d) leaf galls (here, with emerging gall wasp circled in red), and leaf mines: (e) typical leaf mine and (f) leaf mine with miner inside circled in red. We found other herbivores, such as aphids and caterpillars, pressed with plant specimens, but these were rarer. Chewing damage is typically made by caterpillars and beetles; leaf mines are made by flies, beetles, and mites; stippling is made by leafhoppers and other species that remove cell contents from leaves; leaf galls are typically made by gall wasps.

**Fig. 3.**
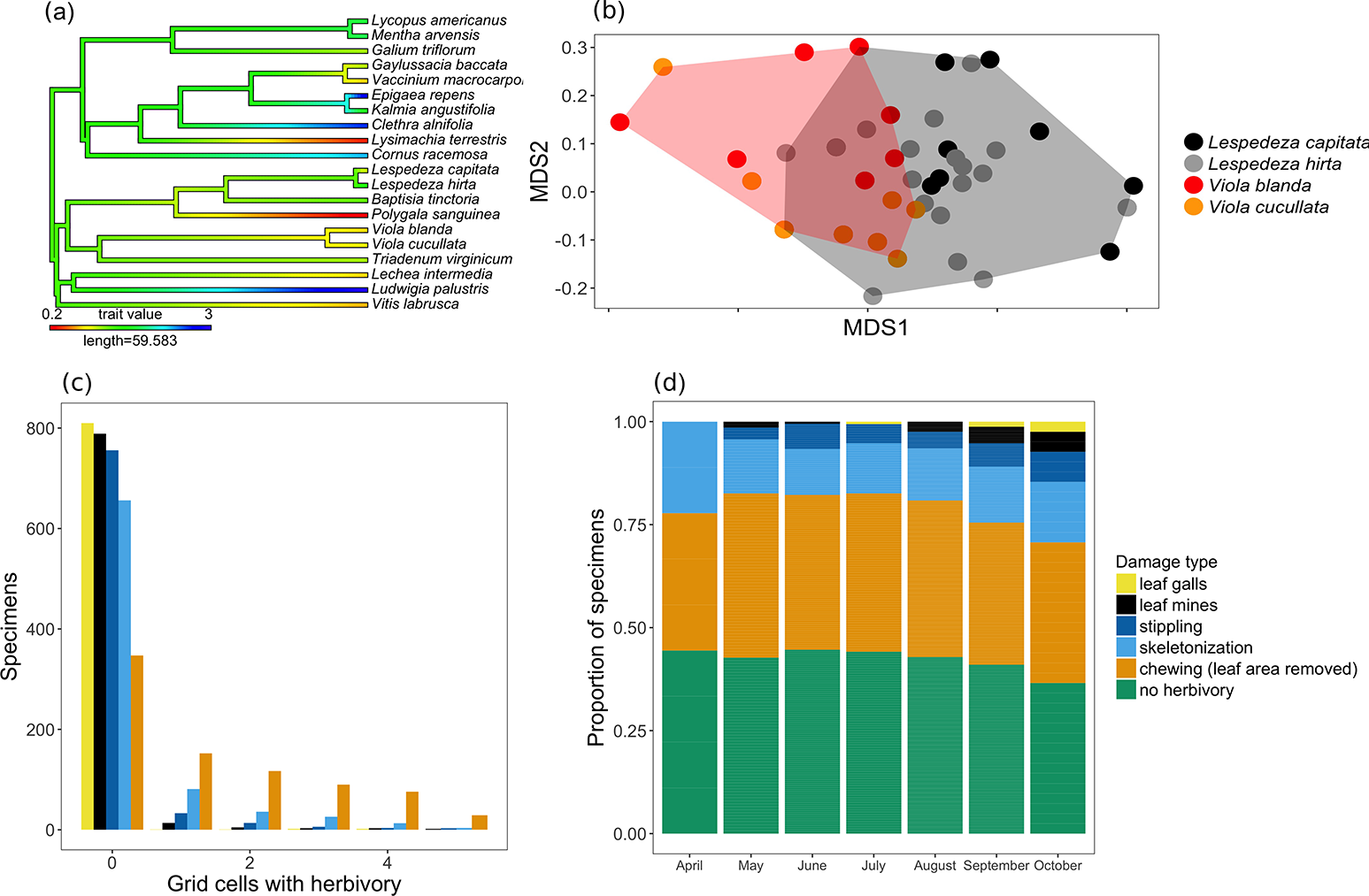
Herbivory recorded on herbarium specimens of New England. We examined herbarium specimens from 20 species from the northeastern US for various types of herbivory (see Appendix S1) and contrasted observations to expectations from the literature. We found that (a) species that are closely related have similar herbivory (Blomberg’s *K*= 0.4), (b) composition of damage types on herbarium specimens is more similar within than between species and genera, illustrated here by two genera Viola and Lespedeza (PERMANOVA, *F*19,507=7.05, *p*<0.001). Herbivory within genera clusters more tightly than between genera, as shown by the general separation of polygons. Similarly, herbivory composition is significantly different between species within genera, shown here as partial overlap of points colored by species. (c) Herbivory on herbarium specimens is highly skewed, and (d) herbivory increased as growing seasons progressed. Chewing damage is shown in (a) and (d) because it was the most prevalent type of herbivory (as shown in c). Detailed statistics are available in Appendix 1.

Collector bias toward intact specimens makes it difficult to infer the absolute magnitude of herbivory from herbarium specimens. However, cross-validation with field data might allow calibration of herbivory estimated from collections and allow researchers to quantify and correct for under-estimates. Another limitation of these data is that we cannot infer herbivory rates, i.e., herbivory per day or month, without data on the timing of leaf-out, though degree-day data may serve as a proxy and thus facilitate these estimates and associated inferences.

### Coarse, incomplete, or inaccurate metadata

In some cases, specimens have been misidentified or taxonomic arrangements revised, but herbarium records have not kept pace with changes. New databases and bioinformatics tools now allow rapid taxonomic name synonymisation (e.g., the taxize R-package, [https://cran.r-project.org/web/packages/taxize/index.html]). Judicious choices of exemplar species can also help researchers avoid problematic taxa, as can sampling within herbaria with well-curated metadata. However, in some cases, reviewing each specimen individually is necessary, and herbarium specimens are themselves the key resource for resolving species identities. In contrast to historical observations, for which it is usually impossible to verify species identifications, herbarium specimens can always be revisited. However, numerous groups likely remain poorly described in herbaria, and despite rapid efforts to mobilize collections online, the resolution of digital images may not be sufficient to provide definitive identification, especially in taxonomic groups with many closely related and morphological similar species, such as in the graminoids.

Another hurdle in using herbarium specimens, especially for ecological applications, is that locations and dates associated with specimens are sometimes absent or incomplete. Label information on specimen locations can be at coarse geographical scales (e.g., at the county level in the US), or is not provided, especially for specimens collected long ago. This is further complicated by the fact that such specimens may be inaccurately georeferenced, which can propagate through subsequent analyses especially if climate data is linked to such data (Park & Davis 2017). Because we cannot go back in time and collect more fine-scale location data, its absence may prevent researchers from using specimens to address certain hypotheses. However, centralized databases, such as Integrated Digitized Biocollections ([https://www.idigbio.org/]), now allow researchers to assess the availability and resolution of data across many herbaria, and thus evaluate whether there is sufficient information to address a hypothesis of interest before investing time in what could be wasted research effort.

### Effects of preservation

Preservation can degrade certain data sources, such as DNA, plant structure, and plant coloration. Technological advances have already helped overcome some of these limitations and may do so to an even greater extent in the future. For example, DNA quality varies considerably among herbarium specimens, but it is already possible to amplify very low, fragmented concentrations of DNA from specimens allowing population and phylogenetic studies using ancient DNA (Särkinen *et al.*, 2012; Applequist & Campbell, 2014). Genomic data may eventually allow us to link genes to key plant traits, thus providing a means to characterize evolutionary responses to environmental stress, pathogens, or competitors that cannot be measured directly from preserved specimens or that have been lost as specimens have become degraded.

Other problems arise because of accidental damage to specimens, notably flooding and insect pests. For example, the global seedbank in Norway was almost flooded in the spring of 2017 due to melting permafrost. In addition, insect pests readily eat herbarium specimens that are not in sealed cabinets, and even herbaria with sealed cabinets can experience infestations, which may not be identified until hundreds of specimens are damaged. While many herbaria use integrated pest management to monitor and treat these occurrences today, historical specimens are often part of personal collections that were exposed to insects. Physical damage of specimens could affect data quality, notably for measurements of herbivory that occurred before collection.

## NEXT GENERATION HERBARIA

We have discussed how herbarium collections can contribute to global change research, and we have described some simple approaches for sampling and analyzing data from herbaria, helping overcome a key barrier to the use of herbarium specimens in ecology. We suggest consideration of sampling strategies which are used in ecology more generally, the adoption of statistical methods to help account for gaps in coverage, and new computational tools, such as those that are appropriate for sparsely sampled data. We emphasize the importance of cross-validating data extracted from herbarium specimens, and suggest how data reliability can be assessed even when companion data are not available. We also identified gaps in the literature that point to future research opportunities. Here, we detail new frontiers for using data derived from herbaria in global change research.

Herbarium specimens could provide unprecedented amounts of data on fungal responses to global change. While herbaria house millions of fungal specimens, these records are rarely integrated into global change research. This presents opportunities for investigating fungal responses to global change in ways that parallel ongoing efforts using plants. Though examples are sparse, herbaria studies have shown increasing incidence of fungal plant diseases (Antonovics *et al.*, 2003) and effects of climate on fungal phenology (Kauserud *et al.*, 2008; Kauserud *et al.*, 2010; Diez *et al.*, 2013). Researchers could draw methods from these studies to more deeply investigate fungal responses, which will be critical determinants of ecosystem function in the future.

Plant and fungal data from herbaria, such as those described in Tables 1 and 2, could be placed on common axes with data from traditional sources—field observations, experiments, and fossils—to generate more robust predictions of how species will respond to drivers of global change (Davis *et al.*, 2015; Youngsteadt *et al.*, 2015). Because all of these methods introduce biases and have limitations, the most robust inferences would include data from multiple approaches, and herbarium data could be uniquely suited in some cases to inform outstanding debates about global change introduced by more traditional approaches. For example, observations and experiments can yield different results about ecologically important phenomena, such as phenological responses to warming (Wolkovich *et al.*, 2012). Herbarium data may allow researchers to resolve such discrepancies, in part, because specimens can capture both long-term processes, including evolution, and short-term processes, such as plastic responses, that are generally not represented together in either observations or experiments. In addition, herbarium records and experiments could be used together to tease apart mechanisms that drive species responses, and herbarium data could be combined with data from fossils (as other modern data have been compared to fossils, as described in Labandeira & Currano 2013) to contrast effects of current and historical climate change by placing temperatures across epochs and recent time on the same axes. While patterns and dynamics might differ due to radically different timescales of these data, such differences might themselves reveal important insights into universal drivers of and responses to global change (e.g., increased CO2 levels and warming).

Leveraging the potential of herbarium data will require advances that allow researchers access to “big data” that span the full range of spatial, temporal, and taxonomic information contained within herbaria and that will require new computational tools to explore. Museum specimen records and images are now more accessible than ever thanks to large scale digitization efforts that have created centralized repositories of these data (e.g., Integrated Digitized Biocollections, Australia’s Virtual Herbarium, Museum National d’ Histoire Naturelle, Paris), although much data remain dark. Extracting ecologically meaningful data from digitized specimens presents additional challenges, notably in the time and resources required. The rapid growth of citizen science, in which the public aid in data collection efforts, has provided one way forward. Recent collaborations between biologists and computer scientists present new opportunities (Willis *et al.*, 2017b) and have allowed for the development of crowdsourcing image annotation tools (e.g., *CrowdCurio*, [https://www.crowdcurio.com/]) to extract phenological data from digitized herbarium specimens. These tools are already being adopted and have enormous power for leveraging herbarium data for climate change research, and preliminary studies suggest they can generate highly usable data despite inherent crowdsourcing error (Williams *et al.*, In Review). Similarly, computer algorithms for analyzing digital images could provide an alternative approach for quantifying traits, such as leaf morphology (Corney *et al.*, 2012a; Corney *et al.*, 2012b; Unger *et al.*, 2016; Wilf *et al.*, 2016) and offer the opportunity to rapidly collect data across large numbers of specimens.

Plant and fungal collections continue to grow, but the goals of herbaria frequently remain unaligned with their ecological utility. There is a need to better integrate current research demands into collecting methods and collection management. Therefore, we suggest a shift in how herbaria operate to help maximize contributions of herbaria to global change research and to center herbaria as key repositories of ecological data. We propose several key strategies to move towards this goal, the last three of which are potentially the most resource intensive:

1) **Organize specimens by evolutionary history (phylogeny) and spatial location.** Collections are most accessible when they are organized by taxonomy and specimen location because researchers almost always sample within clades and areas. Following a more phylogenetically oriented classification and ordering rather than outdated classifications that do not align with current understanding of phylogeny might facilitate ease of usage (see: [http://www.bvaenviro.com/Public/Angiosperm%20Phylogeny%20Group%20III/APG%20III%20linear%20list.pdf]).
2) **Digitize and generate publicly accessible online databases.** While this is not a novel suggestion, we include it to emphasize the importance of publicly available databases and specimen images for supporting ecological research efforts and to acknowledge that the digital tools necessary for making such data available are still evolving. In many cases, research questions do not initially require physical specimens, and online specimen information facilitates assessment of project feasibility, research planning, and sampling. Online databases and images can also serve as sources of preliminary data that can be used in grant applications.
3) **Retain or, at minimum, digitize damaged specimens.** Curators sometimes dispose of specimens that were damaged by insects or pathogens before they were collected. They do this for good reasons; herbaria have limited space and specimen storage comes at a cost. However, damaged specimens contain valuable information on locations and about species interactions. For suggestions on sterile ecological vouchers see below.
4) **Collect and preserve tissue for future molecular and chemical analyses.** Curators at many herbaria now collect and preserve tissue samples, in addition to pressed and dried specimens, explicitly for future molecular analyses. These samples can be dried and stored with silica gel and preserved at room temperature or, more ideally, cryobanked for future extraction. Such infrastructure is also invaluable for preserving RNA, which is essential for investigating gene expression but is often degraded rapidly. These materials are best frozen as soon as possible but field fixatives are now available to reduce the burden of collecting such samples. Recent efforts in tissue banking (see, e.g., [https://frozenark.org/], [http://www.ggbn.org/ggbn_portal/]) provide a useful guide on how such approaches could be implemented.
5) **Add sampling information to specimen labels.** Specimens are most useful when labels include metadata, which are recorded regularly as part of basic collecting protocol, i.e., date, geolocation, species. We suggest that the type of sampling and habitat should also be indicated as standard metadata. For example, one system would be to indicate sampling as targeted or opportunistic, and if targeted, to indicate the intention of the collection, e.g., to document galls on the specimen or to get a collection from a particular area. Indication of a specimen’s immediate and adjacent habitats, e.g., along a roadside, in an urban or transformed environment, or in a forest, would also help ecologists to determine if specimens were collected in appropriate ways for particular studies.
6) **Develop protocols for storing ecological vouchers.** Traditionally, herbaria have focused on maximizing number and morphological diversity within species across their ranges. However, because of increasing interest in historical ecology, there is a need to prioritize specimens that represent ecological effects (Baker *et al.*, 2017). Space is an ongoing issue in herbaria, and ecological vouchers are especially challenging to store, as specimens from a single study can number in the thousands. We suggest that herbaria develop new protocols that allow ecologists to deposit whole digital specimens and small physical samples from plants in their studies. One example might be to reduce these collections in their physical size and to minimally retain sufficient tissue for subsequent DNA extraction. In concert with rapidly developing DNA barcoding methods (see also below), this approach would facilitate the reconstruction of historical patterns of plant community assembly.
7) **Resample areas and clades for which there are strong historical collections.** Many herbaria have collections that are temporally biased toward the 19^th^ and 20^th^ centuries, and few specimens are available from the last 30 years. However, the intensification of global extinction drivers has largely occurred over the past few decades. We suggest that herbaria look to resample areas and clades with strong historical collections, particularly in cases where this objective can be incorporated into planned collection efforts and associated regional/geographic priorities. New collections would allow researchers to make contrasts before and after significant global change. Such efforts are common, and well organized among local amateur naturalist groups.
8) **Create barcode libraries.** As molecular sequencing costs have declined, it is now more reasonable to create digital repositories of genetic data using species-specific markers, allowing samples to be matched to collections using both molecules and morphology. The former may be especially valuable when plant samples lack key identifying traits, such as flowers or leaves. Although controversial, DNA barcoding efforts, such as the Barcode of Life ([http://www.barcodeoflife.org/]), which now holds over 5 million barcode sequences, illustrate the potential of such approaches.

## CONCLUSIONS

Data from herbaria are increasingly being incorporated in to global change research. Researchers are developing creative new methods to understand how diverse factors affect plants, fungi, and their associates. These methods include using herbarium specimens as occurrence and phenological records and as sources of DNA, physiology, and morphology. Data derived from herbaria have wide breadth across space, time, and the tree of life. Several topics of broad interest in global change biology and to which herbaria can contribute remain underexplored but show great promise. Exciting applications include the use of herbaria as blueprints for restoration, signatures of physiological change, and records of changing species interactions. Collections of all kinds are threatened by declines in financial support. The future of herbaria will, in part, depend on their ability to adapt to current research demands and funding priorities. Here, we have emphasized applications to global change research, but a broader dialogue is needed to maximize collection utility across other disciplines. We should recognize that the value of such collections may only become apparent in the future. We must therefore maximize the current use of collections while continuing high standards of preservation to benefit future generations.

## ACKNOWLEDGEMENTS

We thank Barbara Hanrahan, Anthony Brach, and Daniel Park for their knowledge of and assistance within the Harvard University Herbaria. We thank Charlie Willis for providing expert advice and data from Willis *et al.* (2008). We thank Matt Bertone, David Boufford, and Charley Eiseman for their natural history expertise.

## Appendix S1. Detailed sampling methods for scoring herbivory on herbarium specimens

### Herbivory data collection

We chose focal species by selecting from the list of native species included in Willis *et al.*, 2008. We filtered this list for species with over 15 specimens collected between 1900 and 1920, 1920 and 1940, 1940 and 1960 and deposited at the Harvard University Herbaria. We randomly chose 10 native rosids and 10 native asterids from this filtered database to ensure phylogenetic breadth in our survey. We chose only native species to avoid any effects of recent introduction on herbivory. We then randomly selected 40 specimens from each species collected between 1900 and 1960 from Connecticut, Massachusetts, and Rhode Island and preserved in the Harvard University Herbaria collections. We chose these states to minimize spatial variation in the data and these years to exclude impacts of recent global warming, which became pronounced in the 1970s (IPCC 2014).

To quantify herbivory, we randomly subsampled five cells within a 40.64 by 25.40-cm grid overlaid on each specimen. If a cell did not include leaf material we selected another cell. We marked each grid cell for presence/absence of each type of herbivory, e.g., leaf mines, galls, chewing damage.

We encountered several challenges when quantifying herbivory. Historically, many herbarium collections were previously kept in collectors’ homes where they were not protected from insects. Insects can also infest tightly sealed cabinets where herbarium specimens are currently stored. We were only interested in herbivory that happened outdoors. Therefore, we developed methods for distinguishing indoor and outdoor herbivory. We determined that outdoor herbivory is generally distinct from indoor herbivory, in that live plants form toughened, necrotic tissue around the wounds made by herbivores. However, this tissue is more visible on some species than others. Therefore, certain species may not be appropriate for herbarium studies focused on herbivory. In this study, there were several cases in which individual specimens were damaged, and we could not distinguish indoor from outdoor herbivory. When we encountered this issue, we randomly selected another specimen to replace the specimen with ambiguous damage.

### Statistical analyses

First, we examined the relationships between herbivore damage and host plants. We expected significant variation in herbivory between plant species, because plants have evolved to various extents to defend against different herbivores (Bale *et al.*, 2002). We also expected that closely related species would have similar levels of damage, because they are likely to share similar anti-herbivory traits, including defense and tolerance (Pilson 2000; Agrawal & Fishbein 2008; Futuyma & Agrawal 2009; Fornoni 2011), though we note that few studies explore plant tolerance in phylogenetically explicit frameworks. Consistent with these expectations, we demonstrated that there was (1) significant variation in herbivory between plant species (GLMER; *X*^2^_1,19_= 855.50, *P*<0.0001), focusing on chewing damage, which was by far the most prevalent type of herbivory (Fig. 3c), and (2) evidence suggesting close relatives had similar herbivory (Fig. 3a; phylogenetic signal: Blomberg’s *K*= 0.40), although low sample size (*n* = 20) limited our ability to test for statistical significance. We then evaluated the prediction that damage composition would cluster by plant species. Because herbivory is frequently host-plant specific, the types of herbivory on plant species, e.g., chewing, skeletonization, leaf mines, and the relative amounts of herbivory types, should be more similar among individuals within species than between species. We tested this expectation by building a dissimilarity matrix using the abundances of each damage type on each specimen and evaluating the distances between samples among individuals within vs between species. As expected, damage types cluster within species and genera (Fig. 3b), and centroids within the dissimilarity matrix were significantly different among species (PERMANOVA: *F*19,507=7.05, *p*<0.001).

Second, we examined the statistical distribution of herbivory and the accumulation of damage through time. Because plants vary in their defenses against herbivory, and insect attacks can be highly localized, herbivory data tend to have medians of zero, a pattern seen frequently in studies on living plants (Turcotte *et al.*, 2014). In addition, we would expect that as growing seasons progress, the diversity of damage types found on specimens should increase, as the leaves are exposed to herbivores for longer, and as galls and leaf mines grow and become detectable. We found support for both predictions in our data. Most observations showed no evidence of herbivory with a few observations showing high damage (Fig. 3c), but damage diversity, calculated as the number of types of damage per specimen, was higher on specimens collected later in the year (Fig. 3d; GLMER: Wald *Z*=4.06, *p*<0.0001).

Last, we contrasted our estimates of herbivory from herbarium specimens to a database of herbivory collated by Turcotte *et al*, (2014). We found five matching genera between datasets and demonstrated that, across these genera, herbivory was strongly correlated (Pearson correlation: *r* = 0.71).

